# Interaction of lncRNA LENT with DHX36 regulates translation and suppresses autophagy in melanoma

**DOI:** 10.1101/2025.02.03.636228

**Authors:** Alexandre Haller, Giovanni Gambi, Mattia D’Agostino, Guillaume Davidson, Antonin Lallement, Gabrielle Mengus, Chadia Nahy, Nadia Messaddeq, Guillaume Bec, Angelita Simonetti, Eric Ennifar, Irwin Davidson

## Abstract

The melanocyte lineage determining Microphthalmia-associated transcription factor (MITF) drives proliferation and survival of melanocytic melanoma cells through regulation of both coding genes and long non-coding RNAs (LncRNAs). Here we characterize LINC00520 (hereafter called <u>L</u>ncRNA <u>ENhancer</u> of <u>T</u>ranslation, LENT) regulated by MITF and strongly expressed in melanocytic melanoma cells. LENT is essential for proliferation and survival of cultured melanocytic melanoma cells and xenograft tumours. LENT interacts with the G4 quadruplex resolvase DHX36 and both associate with the ribosome in the 80S and light polysome fractions. LENT modulates DHX36 association with a collection of mRNAs regulating their engagement with polysomes and fine-tuning their subsequent translation. These mRNAs encode proteins involved in endoplasmic reticulum (ER) and mitochondrial homeostasis as well as autophagy. Consequently, LENT silencing leads to extensive autophagy and mitophagy, compromised oxidative metabolic capacity accompanied by an accumulation and mis-localization of mitochondrial proteins leading to proteotoxic stress and apoptosis. The LENT-DHX36 axis therefore fine-tunes translation of proteins involved in ER and mitochondrial homeostasis suppressing autophagy and promoting survival and proliferation of melanoma cells.

## Introduction

Melanoma tumours are notoriously heterogeneous, with melanoma cells adopting multiple cell states with differing proliferative, invasive and stem cell capacities (1–7). Intra-tumour heterogeneity is a major determinant of therapeutic resistance with mesenchymal-type cells playing a critical role in targeted and immune checkpoint resistance (4, 7). The transcription programs associated with the different cell states are driven by of a host of transcription factors, with the lineage-defining MITF (Microphthalmia-associated transcription factor) and SOX10 driving the more differentiated melanocytic cell state, while AP1, TEAD, PRRX1 and TCF4 drive the undifferentiated mesenchymal state (6–11). Other intermediate states have been defined such as the neural crest stem cell (NCSC)-like state that plays a key role in minimal residual disease and the emergence of drug resistant populations (3, 12).

In melanocytic melanoma cells, MITF and SOX10 bind together at cis-regulatory elements to promote expression of genes driving proliferation, survival and oxidative metabolism (8, 9). While, these factors regulate multiple coding genes they also regulate expression of long non-coding (lnc)RNAs such as the melanoma-specific lncRNA SAMMSON (LINC01212) essential for melanoma cell proliferation and survival through coordinating mitochondrial and cytoplasmic translation (13, 14). SAMMSON silencing induced mitochondrial precursor overaccumulation stress (mPOS), a form of proteotoxic stress, resulting in melanoma cell death. SOX10 also regulates the melanoma-specific lncRNA LENOX (LINC00518) that interacts with the small GTPase RAP2C promoting its interaction with DRP1 and impairing mitochondrial fission through enhanced DRP1 S637 phosphorylation (15). LENOX potentiates oxidative phosphorylation metabolism to promote melanoma cell survival and resistance to MAP kinase inhibitors (16).

Here we characterize LINC00520 (hereafter called <u>L</u>ncRNA <u>ENhancer</u> of <u>T</u>ranslation, LENT) strongly expressed in melanocytic melanoma cells under the regulation of MITF and essential for proliferation and survival *in vitro* and *in vivo*. LENT interacts with the G4 quadruplex resolvase DHX36 and both associate with the 80S and light polysome fractions. LENT modulates DHX36 association with a collection of mRNAs regulating their engagement in polysomes and their subsequent translation. LENT coordinately regulates engagement with the light polysomes of mRNAs encoding proteins enriched in endoplasmic reticulum (ER) and lysosome homeostasis as well as autophagy and mitophagy. Consequently, LENT silencing leads to extensive autophagy/mitophagy, compromised OxPhos capacity and accumulation and mis-localization of mitochondrial proteins leading to proteotoxic stress and apoptosis. Our results are consistent with a model where LENT fine-tunes translation of proteins involved in ER, lysosomal and mitochondrial homeostasis by modulating the ability of ribosome-associated DHX36 to unwind G4 structures in their mRNAs and their engagement with polysomes. LENT, LENOX and SAMMSON hence constitute a set of melanoma-expressed lncRNAs that act coordinately to fine-tune translation and/or mitochondrial activity and promote melanoma cell proliferation and survival.

## Results

### LENT is expressed in melanocytic melanoma cells and associated with poor patient outcome

Integration of MITF and SOX10 ChIP-seq data with RNA-seq following MITF silencing in 501Mel melanocytic melanoma cells (9) identified LENT (LINC00520) as a lncRNA directly and positively regulated by MITF. LENT expression was reduced upon silencing of MITF or its cofactor BRG1 and the corresponding locus displayed several MITF bound sites associated with BRG1 and marked by H3K27ac in 501Mel cells (**Fig. S1A**). LENT expression was low in normal tissues (0.854 Log2 normalized counts) with highest expression in the oesophagus mucosa and stomach in the GTEX database (**Fig. S1B** and data not shown). Expression was highest in cutaneous melanoma (SKCM, 6.189 Log2 normalized counts), compared to other cancer types (1.490) including uveal melanoma (UVM, 1.732) (**Fig. S1B**). Expression was also upregulated in primary melanoma compared to benign nevi and normal tissues, (**Fig. S1C**). Thus, LENT expression was negligible in normal tissues and upregulated more than 60-fold in cutaneous melanoma.

Analyses of scRNA-seq data from human melanoma xenografts (3) showed that LENT was widely expressed except in NCSC and mesenchymal cells (**Fig. S1D**), while in scRNA-seq data from human melanoma patients (7), it was also broadly expressed except in mesenchymal cells and was strongest in the hypoxia-stress cell cluster (**Fig. S1E**). Preferential LENT expression in melanocytic type cells was confirmed by RT-qPCR analyses of a collection of melanoma cell lines (**Fig. S1F**). Cytoplasmic LENT localization in MITF expressing melanoma cells could be directly observed using RNA-scope on human melanoma patient sections, whereas its expression was low in normal melanocytes (**Fig. S1G**). RNA-scope also showed a predominantly cytoplasmic localization in cultured melanoma cells, whereas no signal was seen in Hela cells (**Fig. S1H**). Further analyses showed the 432 bp isoform 5 as the most abundant in 501Mel cells and also in melanoma patients (**Fig. S2A-C**). LENT is therefore a cytoplasmic melanoma-enriched lncRNA most abundant in melanocytic MITF-expressing cells.

To determine whether LENT expression correlated with patient outcome, we divided the TCGA SKCM dataset into primary and metastatic samples, performed unsupervised clustering of the transcriptome data from each and GSEA analyses of differentially expressed genes to define the signatures of each cluster. In primary melanoma, LENT was co-expressed with MITF and SOX10 in cells defined by an oxidative phosphorylation (OxPhos) and cell cycle signature typical of melanocytic MITF-expressing cells, but was strongly reduced in mesenchymal (designated as EMT, epithelial to mesenchymal transition) cells expressing markers such as PRRX1 (**Fig. S3A)**. In contrast, as previously described (15) LENOX displayed a broader expression pattern being expressed also in EMT cells.

In metastatic melanoma, LENT again was strongest expressed in the MITF-SOX10 expressing OxPhos/cell cycle cells, but reduced in cells with EMT signatures (**Fig. S3B**). In primary melanoma, high LENT expression was associated with better survival whereas in metastatic samples high LENT expression was associated with poorer outcome (**Fig. S3C-D**). These observations are in agreement with and extend previously published analyses (17) and in line with the idea that low LENT-expressing mesenchymal cells promote metastases of primary melanoma (6), whereas LENT-expressing cells with OxPhos and cell cycle signatures associate with poorer survival in metastatic samples, hence accounting for the differential association of LENT expression with survival.

### LENT cooperates with LENOX and SAMMSON to promote melanoma cell proliferation and survival

To address the function of LENT in melanoma cells, we silenced it’s expression by CRISPR interference (CRISPRi) using dCAS9-KAP1, transfection of locked nucleic acid GapmeR antisense oligonucleotides (ASO) or by Doxycycline (Dox)-inducible expression of LENT-targeting shRNA. CRISPRi silencing in 501Mel cells with LENT promoter-targeting sgRNAs that potently reduced its expression resulted in strongly reduced colony forming capacity (**Fig. 1A-B**). Transfection of melanoma cells with different phenotypes and driver mutations, with 2 independent ASOs that reduced LENT expression by over 80% compared to a non-targeting control (CTR) (**Fig. 1C and Fig. S2 D-E)** and reduced growth of melanocytic, but not mesenchymal melanoma cells nor HEK293T cells that did not express LENT (**Fig. S4A**). ASO-mediated silencing resulted in strongly reduced cell proliferation (**Fig. 1D**) and a strong increase in cleaved caspase 3-expressing apoptotic cells (**Fig. 1E**) with early and late apoptotic cells observed in flow cytometry (**Fig. S4B**). We also silenced LENT with a stably integrated Dox-inducible shRNA that efficiently reduced 501Mel proliferation (**Figs. S2E and S4C**). In contrast, ectopic Dox-induced expression of LENT isoform 5 stimulated colony formation in melanoma cells, but also in HEK293T cells where it was not normally expressed **(Fig. S4D-E**)

**Figure 1:**
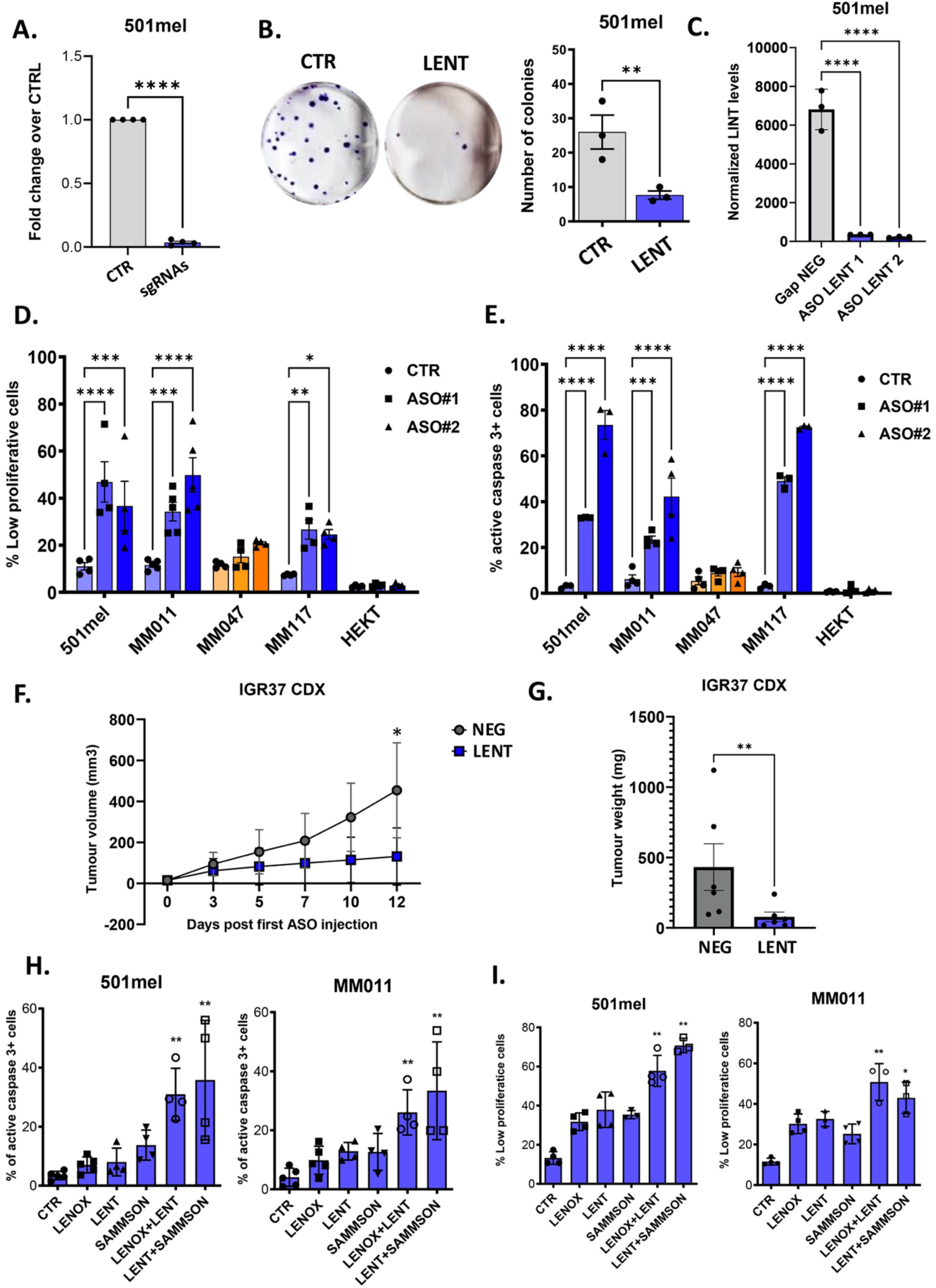
LENT depletion impairs melanoma cell proliferation and survival. **A.** LENT levels measured by RT-qPCR after dCas9-KAP1-mediated LENT silencing compared by paired t-test. **B.** Colony formation assay upon dCas9-KAP1-mediated LENT silencing, compared by one-way ANOVA (Dunnett test). **C.** LENT levels measured by RT-qPCR after ASO-mediated depletion with two independent ASOs. **D-E.** Measurement of slow proliferating or apoptotic cells by flow cytometry upon ASO-mediated LENT depletion compared by one-way ANOVA (Dunnett test). Melanocytic cell lines are represented in blue and mesenchymal cells are colored in orange. **F-G.** Tumour volumes in mice with IGR37 CDX tumours were measured at the indicated number of days following initial injections of LENT-targeting ASO. Tumours were weighed following sacrifice at day 14. Following the first ASO injection. Volumes were compared by two-way ANOVA and weights by Mann-Whitney test. **H-I.** Same measurements as for D-E but after ASO-mediated depletion of the indicated lncRNAs at sub-optimal ASO doses, compared by one-way ANOVA (Dunnett test). *, P < 0.033; **, P < 0.0021; ***, P < 0.0002; ****, P < 0.0001.

To test if LENT silencing could also block xenograft tumour growth, melanocytic IGR37 cells were injected subcutaneously in immunodeficient mice and when tumours reached ≈100 mm^3^, mice were subsequently injected subcutaneously every 2 days with LENT ASO. Compared to untreated controls, injection of LENT ASO reduced LENT expression in tumours and strongly reduced tumour growth and tumour weight (**Fig. 1F-G, and Fig. S4F**).

All 3 targeting strategies as well as gain of function therefore revealed the essential role of LENT in the proliferation and survival of melanocytic melanoma cells in culture and *in vivo* xenografts similar to previous reports (17).

We previously showed that ASO silencing of LENOX and SAMMSON cooperated to induce melanoma cell death (15). To assess if LENT also collaborated with LENOX and SAMMSON, melanocytic 501Mel and MM011 cells were transfected with sub-optimal concentrations of ASO targeting LENT alone or together with LENOX or SAMMSON. Each ASO specifically targeted its cognate lncRNA target without affecting expression of the others with the exception of SAMMSON that was upregulated in the LENT-LENOX targeted MM011 cells (**Fig. S5A**). Compared to LENT, LENOX or SAMMSON alone, a cooperative increase in apoptosis of both lines was observed using the combinations of ASO, and an additive increase in slow proliferation (**Fig. 1H-I)**. LENT silencing also cooperated with MAP Kinase inhibition by the dabrafenib and trametinib combination to eradicate melanoma cells (**Fig. S5B**). MITF and SOX10 therefore coordinately regulate a network of 3 lncRNAs that cooperate to promote melanoma cell survival.

### LENT interacts with the G4 resolvase DHX36

Consistent with the observation that LENT is predominantly cytoplasmic and so less likely to influence transcriptional regulation, RNA-seq from ASO-control or LENT ASO2-silenced cells revealed 97 up-regulated and 82 downregulated transcripts (Log2 fold-change +/-1 p<0,05). LENT silencing did not have a major impact on gene expression (**Fig. S6** and **Dataset S1**) and may therefore mainly act via other cellular processes.

To identify LENT interacting proteins, we performed pulldown from cytoplasmic extracts of 501Mel cells using a tiling array of biotinylated oligonucleotides complementary to LENT or as negative control, the prostate cancer lincRNA PCA3, followed by mass-spectrometry. Compared to several control lncRNAs, LENT was selectively enriched using its cognate oligonucleotides, but not those of the PCA3 control (**Fig. 2A**). Triplicate purifications were performed and LENT-interacting proteins identified by mass-spectrometry. DHX36 was the most enriched protein in the LENT pulldown with no peptides found in the 3 control samples, but an average of 17 in the LENT pulldowns (**Fig. 2B and Dataset S2**). To confirm this interaction, we performed LENT pulldown from native or UV-crosslinked extracts followed by immunoblot. Under both conditions, DHX36 was enriched in the LENT pulldown compared to the PCA3 control, whereas neither the SAMMSON-interacting CARF (14) nor LENOX-interacting RAP2 (15) were enriched (**Fig. 2C**). For further confirmation, we performed LENT pulldown from the HEK293T cells ectopically expressing LENT isoform 5. DHX36 was detected after pulldown from LENT-expressing HEK293T cells, but not from control cells with empty GFP vector (**Fig. 2D**). In the converse experiment, we immunoprecipitated DHX36 from 501Mel cells and found enrichment of LENT relative to the control IgG and to SAMMSON and MALAT1 (**Fig. 2E-F**).

**Figure 2:**
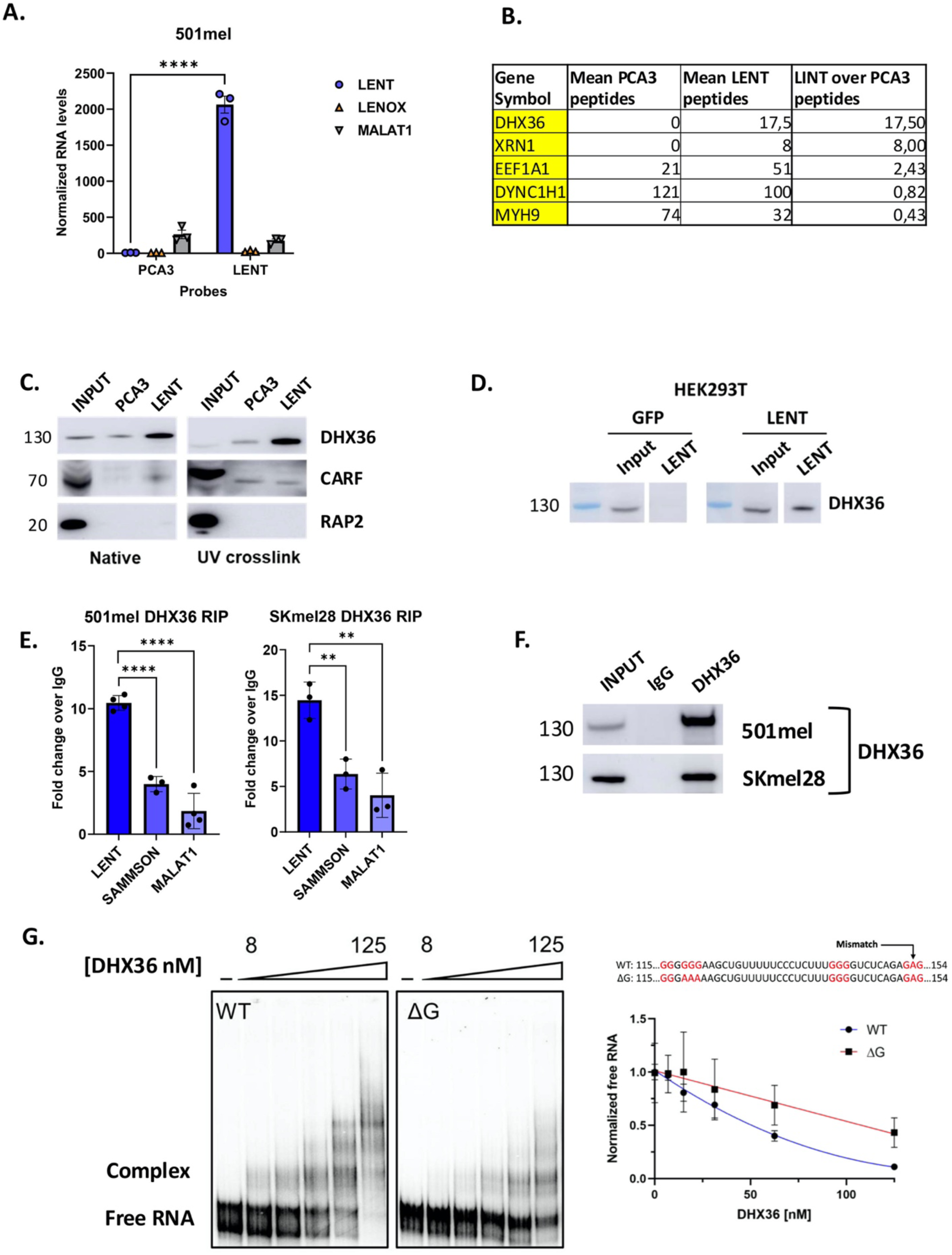
LENT interacts specifically and directly with the G4 resolvase DHX36. **A.** Measurement of lncRNA levels by RT-qPCR after RNA pulldown with control probes (PCA3) or LENT-targeting probes by two-way ANOVA. **B.** LC/LC Mass spectrometry analysis of peptides retrieved after biological triplicate pulldown of PCA3 or LENT. **C.** Immunoblot showing protein enrichment after the indicated RNA pulldowns, in native or UV crosslinked conditions. **D.** Native LENT pulldown upon ectopic expression of GFP or LENT in HEK293T cells. **E.** Enriched lncRNAs analysed by RT-qPCR after DHX36 immunoprecipitation compared with a control IgG by unpaired t-test. **F.** DHX36 immunoprecipitation using a specific antibody showing DHX36 enrichment by immunoblot. **G.** EMSA assay performed with T7 in vitro transcribed WT or mutated LENT in presence of increasing concentrations of purified truncated DHX36. The potential G4-forming structure predicted by PQSfinder in LENT isoform 5 sequence is indicated in red. The mutated LENT sequence used for the EMSA with the three guanines mutated in adenines underlined. *, P < 0.033; **, P < 0.0021; ***, P < 0.0002; ****, P < 0.0001.

To ask if LENT interacts directly with DHX36, we generated and purified recombinant DHX36 in *E.Coli* (**Fig. S7A-B**) and performed electrophoretic mobility shift assay (EMSA) with *in vitro* transcribed LENT isoform 5 RNA (**Fig. 2G**). The presence of increasing amounts of purified DHX36 shifted LENT into slower migrating DHX36-RNA complexes. As DHX36 binds RNA with G4 structures (18), we used the QGRS program (19) that predicted a potential G4 structure in LENT, but with a rather low score. Mutation of three guanines in this sequence decreased complex formation in EMSA, but did not fully abolish the interaction (**Fig. 2G**). Together, these *in cellulo* and *in vitro* experiments revealed a selective and direct interaction of LENT with DHX36 that is partially dependent on a potential G4 forming sequence in LENT.

### LENT modulates association of mRNAs with DHX36

DHX36 unwinds G4 structures in both DNA and RNA and in particular in the 5’-UTR of mRNAs to facilitate their translation (18, 20–23). This observation suggested that LENT may modify DHX36 interactions with mRNAs and their translation in melanoma cells. We therefore investigated the mRNAs associated with DHX36 and determined if their association was modulated by LENT silencing. We performed triplicate DHX36 or control IgG immunoprecipitations (IP) from 501Mel cells expressing control shRNA and the associated mRNAs were sequenced to identify those associated with DHX36 in control conditions. Almost 2000 mRNAs were enriched in the DHX36 IP compared to IgG, whereas 1949 mRNAs were less present in the DHX36 IP compared to control (Log2 fold-change +/-1 p<0,05) (**Fig. 3A** and **Dataset S3**). One of the most enriched was the DHX36 mRNA suggesting DHX36 acts to regulate its own translation in a positive regulatory loop. Ontology analysis showed that DHX36-associated mRNAs were enriched in those encoding proteins involved in mitochondrial function with protein targeting to mitochondrion, mitochondrial calcium ion homeostasis amongst the most enriched terms (**Fig. 3B**). Comparison with RNA-seq data from 501Mel cells showed no correlation between association with DHX36 and expression levels excluding the possibility that we spuriously enriched highly expressed mRNAs in the DXH36 IP (**Fig. S8A**).

**Figure 3:**
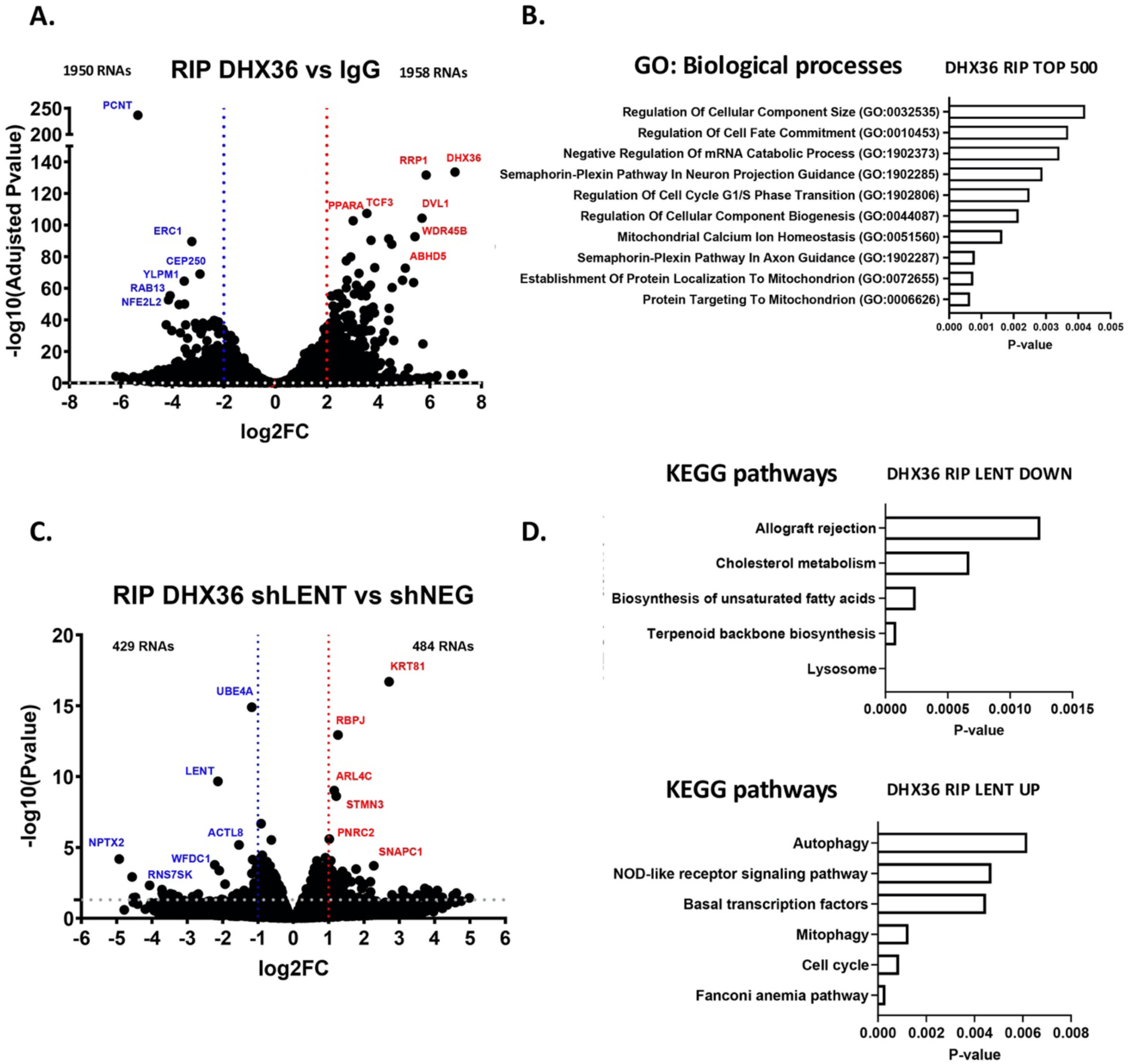
LENT modulates association of RNA with DHX36. **A.** Volcano blot showing RNAs enriched or depleted in the DHX36 IP vs control IgG IP. P-values were derived using the Wald test and adjusted using Benjamini-Hochberg FDR correction. **B.** Gene ontology analysis by EnrichR software of the 500 most enriched RNAs in the DHX36 IP. **C.** Volcano blot showing RNAs enriched or depleted in the DHX36 IP upon shRNA-mediated LENT silencing. P-values were derived using the Wald test. **D.** KEGG pathways by EnrichR of RNAs enriched or depleted in DHX36 IP upon LENT depletion.

It has been reported that mRNAs that associate with DHX36 are enriched in a GG-rich motif with a propensity to form G4 structures [(24) and **Fig S8B**]. This motif was predicted to be present in around 20% of the mRNAs enriched in the control IP, but close to 40% in the DHX36 IP (**Fig. S8C**) and increased to 50-60 % when considering the mRNAs most enriched in the DHX36 IP (**Fig. S8C** and **Dataset S3**). Examination of the DHX36 mRNA sequence with QGRS mapper indeed identified several potential G4-forming sequences including the 5’-GGnGGnGG-3’ motif (**Fig. S8D**) consistent with the observation that it was one of the most enriched mRNAs. We compared mRNAs enriched in the DHX36 IP in 501Mel cells with previously published RNA-seq data from HeLa or HEK293T cells designed to identify G4-containing RNAs (25, 26). Comparing the overlap between the 2 HeLa datasets showed 1832 common mRNAs representing between 46% and 64% of the identified G4-containing mRNAs using different techniques. Comparison with the 501Mel mRNAs enriched by DHX36 IP showed that 900 (46%) were shared with the HeLa and HEK293T datasets, with 341 common to all (**Fig. S8E**). These common mRNAs were enriched in terms associated with transcription and MAP Kinase signaling (**Fig. S8F**).

We also compared the top 500 RNAs in our dataset with the 500 most enriched in a previously published DHX36 PAR-Clip (27) dataset. Among them, only 41 were common between the two datasets, including the mRNA encoding DHX36 (**Fig. S8G**). We observed a clear enrichment for mitochondria-related terms in the 501Mel DHX36 IP (**Fig. S8H**) compared to Sauer et al, characterized by transcription and TGF-B signaling related terms (**Fig. S8I**). This comparison was however potentially confounded by the fact that HeLa and 501Mel melanoma cells have very different gene expression profiles.

We then investigated if LENT silencing modified mRNA interaction with DHX36. We directly compared RNA-seq of the DHX36 IP from the control shRNA with the DHX36 IP from the shLENT cells and identified 484 genes displaying increased association with DHX36 in absence of LENT and 429 with less association (p<0,05) (**Fig. 3C**, and **Dataset S3**). As expected, due to its downregulation by shRNA silencing, LENT was identified as less associated with DHX36. Ontology analyses of mRNAs showing increased DHX36 association revealed enrichment in several process including cell cycle, mitophagy and autophagy, whereas those less associated were enriched in lysosome, metabolic process and allograft rejection (**Fig. 3D)**. Analyses of the mRNAs whose association with DHX36 was affected with the QUADRatlas software showed that a large majority comprised experimentally described and/or predicted G4s (**Fig. S8J**). Together these data define DHX36-associated RNAs in melanoma cells and identify RNAs whose association with DHX36 was positively or negatively modulated by LENT.

### LENT and DHX36 are associated with the ribosome and regulate coordinate engagement of mRNAs encoding proteins involved in ER homeostasis with polysomes

While preparing the DHX36-associated RNAs for sequencing, we noted that the 28S and 18S rRNAs were strongly enriched in the DHX36 IP, but not the control IP (**Fig. 4A**) and therefore used ribo-depletion kits to prepare the libraries for RNA-seq. This observation however strongly suggested that DHX36 was associated with the ribosome. To assess this, we performed polysome profiling of 501Mel cell extracts and analyzed both RNA and protein contents of the fractions. Based on the RNA absorption profile (**Fig. 4B**) and the distribution profiles of EIF4A2 (initiation factor marking the 40S) and RPL36 (component of the large subunit), we designated the 40S, 60S, 80S and polysome fractions. DHX36 showed association with the 60S, 80S and was additionally present in the polysome fractions (**Fig. 4C**). As expected, the control GAPDH mRNA was enriched in the heavier polysome fractions, whereas LENT showed a strong peak in the 80S fraction, but rapidly decreased in the light polysome fractions (**Fig. 4D)**. These observations suggested that LENT may associate with DHX36 on the 80S and light polysome fractions.

**Figure 4:**
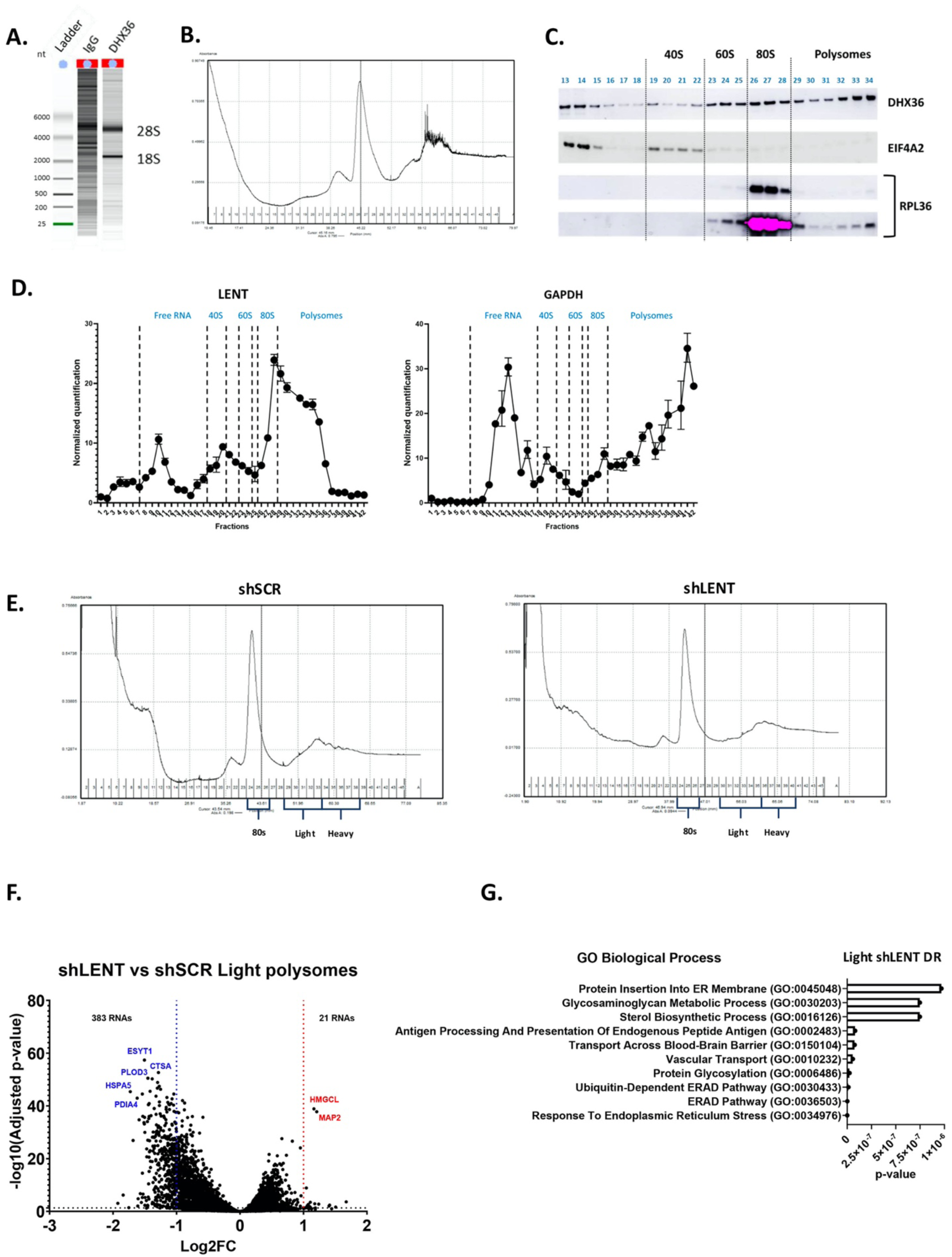
LENT regulates polysome engagement of a subset of mRNAs. **A.** Bio-analyser analyses of precipitated RNA show strong enrichment of 18S and 28S rRNAs in the DHX36 IP but not the control IgG IP. **B.** Sucrose gradient separation of ribosomes with rRNAs measured by UV-light spectrometry. **C.** Immunoblots for DHX36, the initiation factor EIF4A2 and the large ribosomal subunit RPL36 after polysome separation. EIF4A2 is enriched in the 40S and RPL36 in the 80S and heavier polysome fractions. **D.** Presence of LENT or GAPDH mRNA in fractionated ribosomes as measured by RT-qPCR. **E.** Representative sucrose gradient separation of ribosomes with rRNAs measured by UV-light spectrometry. Fractions pooled for the 80S, LP and HP are indicated. **F**. Volcano blot showing RNAs enriched or depleted in the LP fraction from control or shLENT cells. **G.** Gene ontology analysis by EnrichR software of the 383 depleted RNAs in the shLENT LP fraction.

To investigate if the LENT-DHX36 axis regulated mRNA association with ribosomes, we prepared biological triplicate polysome fractions from shControl or shLENT-silenced 501Mel cells. We pooled RNA from four fractions representing the 80S, light or heavy polysome components from each replicate and assessed their composition by RNA-seq (**Fig. 4E** and **Dataset S4**). Few RNAs showed differential presence in the heavy polysome (HP) fractions from the control or LENT silenced cells, whereas 189 and 246 mRNAs were depleted or enriched, respectively in the 80S fraction (Log2 fold-change +/-1 p<0,05) (**Fig S9A-B** and **Dataset S4**). However, the most striking effect was seen in the light polysome (LP) fractions, where 383 mRNAs were depleted in the LENT silenced cells, while only 21 were enriched (**Fig. 4F**).

Ontology analyses of the 383 mRNAs depleted in the LP fractions revealed their strong enrichment in several pathways pertaining to endoplasmic reticulum (ER) homeostasis such as ER stress, ER-associated protein degradation (ERAD), and protein glycosylation (**Fig. 4G** and **Dataset S5**). KEGG ontology analyses gave comparable results, but further revealed enrichment in lysosome function (**Dataset S5**). Key components of the ER stress/ERAD pathways such as the E3 ligase SYNV1, the HSPA5 chaperone, and the PDIA3, -4, and -6 enzymes were all significant depleted in the LP fractions, with PDIA encoding mRNAs also depleted in the 80S fraction (**Fig. 5A** and **Fig. S9C**). The mRNA encoding WFS1 involved in ER Ca2+ transport was also depleted in the LP fractions, along with those encoding the NOMO1, -2 and -3 proteins and other components of the multi-pass translocon complex **(Fig. 5B and Dataset S5)**. Similarly, mRNAs encoding the DPAGT1 and GALNT2, 7 and 12 enzymes involved in N-linked or O-linked protein glycosylation, respectively, were all depleted in the LP fractions (**Dataset S5)**. Related to the above, mRNAs encoding the MHC class 1 HLA-A, -B and -C antigens as well as the TAP1, TAP2 and CALR proteins involved in their transport and antigen presentation were depleted in the LP fractions (**Fig. S9D-E and Dataset S4-S5)**. Overall, these transcripts showed no bias towards low expression compared to the overall transcriptome, but rather were well expressed. Furthermore, they displayed no bias in length compared to the overall transcripts of the LP fraction, although as may be expected transcripts in the LP were significantly longer than those in the 80S fraction (**Fig. S10A-B)**. These results indicated that engagement in LPs of mRNAs encoding key components of many processes associated with normal ER homeostasis and/or intracellular protein transport was coordinately regulated by LENT.

**Figure 5:**
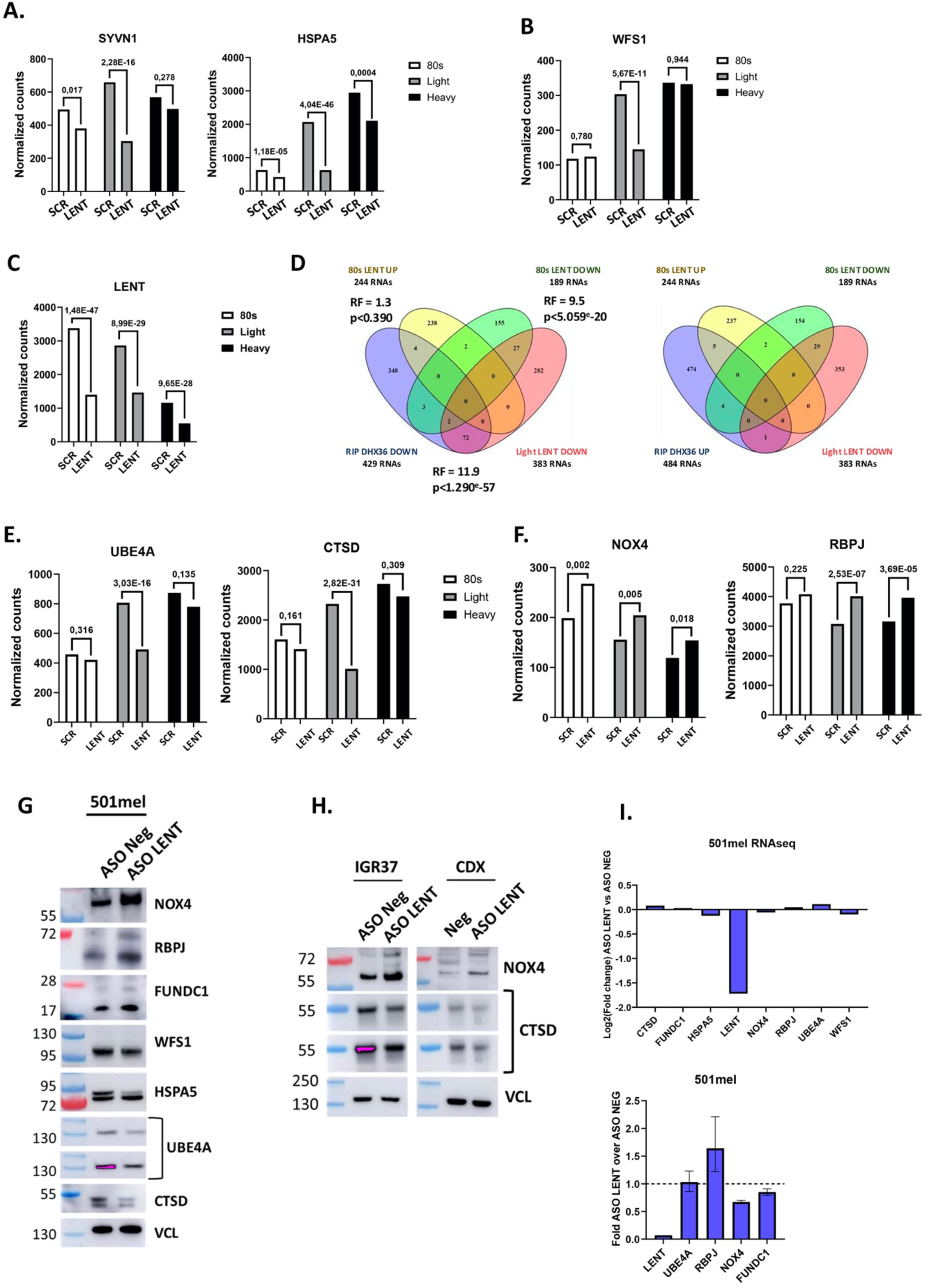
LENT fine-tunes translation of mRNAs differentially associated with polysomes. **A-C.** RNA-seq data showing the representation of LENT in the 80S, LP and HP fractions. The normalized number of reads are shown along with the adjusted p-value between the control and shLENT conditions. **D**. Venn diagrams comparing RNAs modulated in the DHX36 RIP in presence or absence of LENT silencing with those differentially present in the 80S and LP fractions. Significant Representation Factors are shown. **E-F**. RNA-seq data showing the representation of the indicated RNAs in the 80S, LP and HP fractions. The normalized number of reads are shown along with the adjusted p-value between the control and shLENT conditions. **G.** Immunoblots of the indicated proteins 48 hours following ASO-mediated LENT silencing in 501Mel cells. **H.** Immunoblots of the indicated proteins 48 hours following ASO-mediated LENT silencing in the IGR37 cell line or extracts from IGR37 CDX tumours from control or LENT ASO-injected mice. **I.** Expression of the indicated RNAs in ASO control or ASO LENT transfected cells represented as fold-change in RNA-seq (upper panel) or RT-qPCR (lower panel).

We interrogated the polysome RNA-seq to determine if the mRNAs whose association with DHX36 was positively or negative regulated by LENT were also differentially engaged with the polysome fractions. Depletion of LENT was clearly seen in all fractions (**Fig. 5C**). Of the 383 RNAs depleted in the LP fractions, 74 were also depleted in the DHX36 RIP upon LENT silencing (**Fig. 5D** and **Dataset S4**) representing a highly significant, but incomplete overlap between the 2 experimental approaches. In contrast, almost no RNAs showed discordant regulation, with only a single transcript up in DHX36 RIP and down in the LP fractions. Moreover, a smaller, but significant, overlap (29 transcripts) was seen with the transcripts depleted in the 80S fraction, with only 4 discordant transcripts. Ontology analyses of the 74 common mRNAs revealed their strong enrichment in ER homeostasis, including the above-mentioned HLA proteins, and lysosome function analogous to the 383 LP-depleted transcripts (**Fig. S10C and Dataset S4-S5**).

The volcano plot of the LP fraction RNA-seq (**Fig. 4F**). appeared to show a general bias towards the depleted transcripts. This is not due to the presence of a higher number of reduced transcripts, but rather their stronger adjusted pvalues, not only of the transcripts that pass the cut off value, but also transcripts just below the cut off. To further consolidate this analysis, we reanalyzed the RNA-seq to identify 311 transcripts with fold changes between -0.8 and -1.0 with adjusted pvalues < 0,05 that were enriched in ER and Lysosome ontology terms analogous to the 383 transcripts (**Dataset S5**). Taking these additional transcripts into account, a highly significant overlap of 118 transcripts was observed between those depleted in the LP fraction and those with reduced association with DHX36 in the LENT silenced cells, with only 2 transcripts showing a discordant regulation **(Fig. S10D)**. LP depleted transcripts close to cut off values were hence enriched in analogous ontologies to those that pass the cut off. Similarly, analyses of the RNAs depleted in the 80S fraction using relaxed criteria of Log2 fold-change ≥0,7, but with a more stringent adjusted p-value of <0,01 also showed a strong enrichment in many of the same terms related to ER homeostasis (**Dataset S5**).

Together, these analyses showed the effect of LENT depletion on mRNA-association with both the 80S and LP fractions was neither random nor general, but selectively affected transcripts with specific functions that displayed a highly significant, but incomplete overlap with those whose association with DHX36 was promoted by LENT. We note that these transcripts do not overlap with those whose expression was altered upon LENT silencing.

QUADRatlas analyses of the 383 LP-depleted mRNAs indicated the presence of experimentally defined (342/368) and predicted (217/368) G4 forming sequences (**Fig. S10E**). This was consistent with the idea that the LENT-DHX36 axis regulated their unwinding to facilitate their translation. To test this, we investigated if mRNAs whose association with DHX36 and/or engagement with the LP fractions was modified by LENT silencing were differentially translated. The mRNAs encoding UBE4A and CTSD whose interaction with DHX36 was reduced upon LENT silencing also showed reduced association with 80S, LP and HP fractions, with the strongest reduction in the LP fraction **(Fig. 5E).** UBE4A and CTSD accumulated to lower levels following ASO-mediated LENT silencing, whereas RNA-seq and/or RT-qPCR showed no change in overall abundance of the corresponding mRNAs (**Fig. 5G and I)**. The decreased protein level was therefore most likely due to altered translation.

In contrast, mRNAs encoding the ROS generating enzyme NOX4 known to promote melanoma (28) and RBPJ whose association with DHX36 was increased upon LENT silencing were also increased the 80S, LP and HP fractions and although the fold change was below cutoff, their increased association was statistically significant (**Fig. 5F)**. Protein levels of NOX4, FUNDC1 and RBPJ were increased upon ASO-mediated LENT silencing with again no overall changes in the corresponding mRNA levels (**Fig. 5G and I)**. Furthermore, increased NOX4 and reduced CTSD levels were seen in extracts from IGR37 xenograft tumours treated with LENT ASO (**Fig. 5H)**. Similarly, levels of HSPA5 and WFS1 proteins whose RNAs were less associated with the LP fractions were also decreased despite the fact that we did not detect changes in their association with DHX36 (**Fig. 5G)**. These data showed that LENT modulated interactions of mRNAs with DHX36 and/or the ribosome LP fractions to regulate their translation.

### LENT and DHX36 are enriched at mitochondria

Previous studies showed that DHX36 was predominantly cytoplasmic (22, 27). Immunofluorescence revealed that DHX36 was enriched at cytoplasmic structures in 501Mel, IGR37 and A375 melanoma cells (A375; NCSC-type cells, not expressing LENT) that co-staining with HSP60 identified as mitochondria (**Fig. 6A-B**). Co-staining with HSP60 was less prominent in HeLa cells and while little nuclear staining was seen in 501Mel cells, stronger staining was seen in IGR37. ASO-mediated LENT silencing did not modify DHX36 mitochondrial localization, nor did it affect expression of DHX36 mRNA or protein (**Fig. S11A-C**).

**Figure 6:**
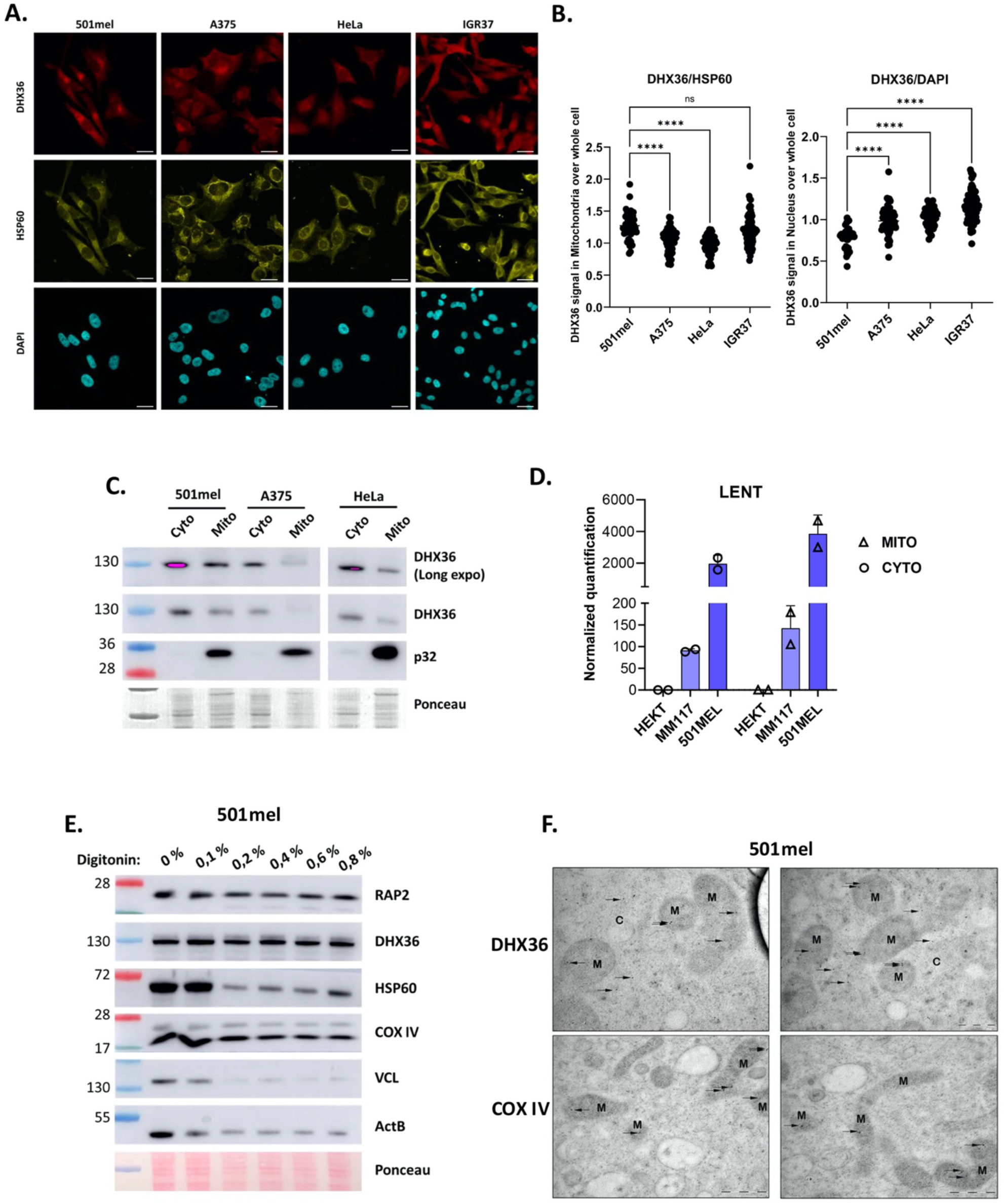
LENT and DHX36 are mainly localized at mitochondria in melanoma cells. **A.** Immunofluorescence by confocal microscopy showing DHX36 localization in different cell lines. HSP60 is used as a mitochondria marker and DAPI to stain the nucleus. Scale bars = 10 µM. **B.** DHX36 signal was quantified in the whole cell and divided by the signal in mitochondria or in the nucleus. Each measured cell is represented by one point and groups compared by one-way ANOVA (Dunnett test). *, P < 0.033; **, P < 0.0021; ***, P < 0.0002; ****, P < 0.0001. **C-D.** DHX36 and LENT levels quantified by immunoblot or RT-qPCR in cytosolic or mitochondrial fractions of different cell lines. **E.** Purified mitochondria were digested with increasing concentrations of digitonin and retained proteins were analyzed by western blot. **F.** Immuno-gold staining coupled with electron microscopy. Representative stained particles for DHX36 or COX IV in the cytoplasm (C) or mitochondria (M) are indicated with arrows. Scale bars are indicated on images.

DHX36 association with mitochondria in 501Mel cells was confirmed by immunoblots of cytoplasmic and mitochondrial fractions (**Fig. 6C**). Similarly, RT-qPCR showed that LENT was also abundant in the mitochondrial fraction from melanoma cells (**Fig. 6D**). We then performed immunoblots of the mitochondrial fraction in presence of increasing quantities of digitonin that was previously used to assess association of proteins with mitochondria (29). While, the control Vinculin (VCL) and Beta-actin (ACTB) proteins were rapidly depleted with increasing digitonin concentration, the mitochondrial protein COX IV was resistant to the highest concentrations (**Fig. 6E**). Both DHX36 and the LENOX-interacting mitochondrial partner RAP2 were also resistant to digitonin showing they were strongly associated with mitochondria. Consistent with this, COX IV and DHX36 showed resistance to tryptic digestion in swelling buffer, whereas VCL and ACTB were sensitive (**Fig. S12A**). To consolidate the idea that DHX36 was located in the mitochondria, we performed immune-gold staining coupled to electron microscopy (EM). Numerous DHX36 particles were detected both in the cytoplasm and within the mitochondria, whereas staining for the COX IV displayed as expected mitochondrial localization (**Fig. 6F**). Hence in melanocytic melanoma cells, a fraction of DHX36 was localized within the mitochondria.

Not only was DHX36 associated with mitochondria, but several of the proteins whose translation it regulated were also associated with mitochondria. Immunostaining showed that UBE4A was strongly enriched at mitochondria in 501Mel and IGR37 cells (**Fig. S12B**). Similarly, while a fraction of NOX4 was present in the nucleus, it was also enriched at mitochondria (**Fig. S12C)**. The transcriptional regulator, RBPJ was mainly nuclear, but a fraction of RBPJ could also be detected at mitochondria (**Fig. S12D**). NOX4, RBPJ and UBE4A were further detected by immunoblot in biochemically purified mitochondria (**Fig. S12E)**. These observations supported the idea that a subset of the proteins regulated by the LENT-DHX36 axis were associated with the mitochondria.

### LENT silencing induces autophagy/mitophagy and proteotoxic stress

The above observations showed that LENT modulated interaction of mRNAs involved in ER-homeostasis, lysosome and autophagy/mitophagy with DHX36 and/or the LP fractions and their subsequent translation. We therefore investigated if LENT silencing impacted these processes. EM showed that LENT-silenced cells were characterized by lower numbers of mitochondria, but an accumulation of numerous autophagosomes not seen in the control shRNA cells (Auto in **Fig. 7A**). Many autophagosomes comprised mitochondria identifiable by their cristae indicating extensive mitophagy. In agreement with this observation, immunoblot with anti-LC3 antibody revealed accumulation of the LC3-II form indicative of autophagy in LENT silenced 501Mel and MM117 cells (**Fig. 7B**). A more modest but detectable LC3-II accumulation was also observed in extracts from LENT ASO-treated IG37 tumours (**Fig. 7C**). Staining of control and LENT silenced cells with both lysotracker and mitotracker showed increased numbers of lysosome-mitochondrial contacts in LENT silenced cells that was further indicative of auto/mitophagy (**Fig. 7D**). Accumulation of autophagosomes was not seen in LENOX silenced cells despite that fact that its silencing impacted mitochondrial homeostasis (**Fig. S13**) (15). Thus, induction of autophagy/mitophagy were major phenotypes of LENT silencing.

**Figure 7:**
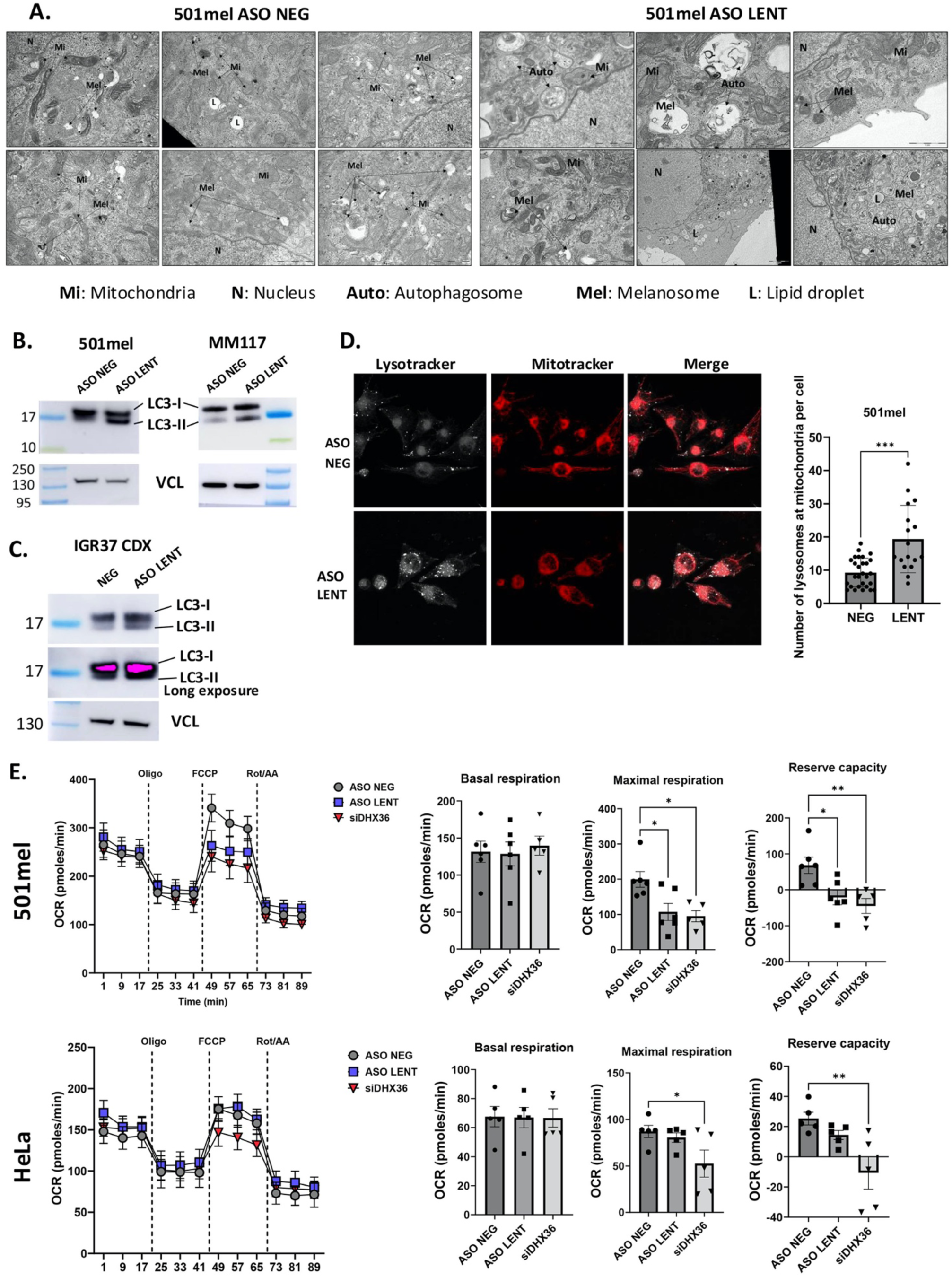
LENT depletion induces mitophagy and reduces oxygen consumption rate. **A.** Transmission electron microscopy of 501Mel cells 48 hours following transfection of control or LENT-targeting ASO. Autophagosomes and degraded melanosomes were observed in LENT silenced cells. Scale bars are indicated on the images. **B.** Quantification of cytosolic LC3B (LC3-I) and lipid-associated LC3B (LC3 II) by western blot in 501Mel or MM117 cells. Vinculin is used as a loading control. **C.** LC3B immunoblot in extracts of IGR37 CDX tumours. **D.** Confocal microscopy of unfixed 501Mel cells stained with lysotracker and mitotracker in control or LENT silenced conditions. The numbers of co-localizing lysosomes and mitochondria was determined and compared between the two conditions by Welch’s test. Scale bars = 10 µM. **E.** Oxygen consumption rate was determined upon LENT or DHX36 silencing. Reserve capacity was obtained by subtracting the maximal capacity with the basal capacity. Comparisons were done by one-way ANOVA (Dunnett test). *, P < 0.033; **, P < 0.0021; ***, P < 0.0002; ****, P < 0.0001.

Given these observations, we asked if mitochondrial function was impacted by profiling the Oxygen Consumption Rate (OCR) using the Agilent SeaHorse. Compared to control ASO, LENT silencing reduced maximal OCR and reserve capacity, but not basal levels in 501Mel cells, whereas no effect was seen in HeLa cells (**Fig. 7E**). DHX36 silencing reduced maximal and reserve capacities in both cell types revealing its more general role in regulating mitochondrial activity (**Fig. 7E**). The mitophagy and impaired mitochondrial function upon LENT silencing led to increased ROS levels and the appearance of ROS-high apoptotic cells (**Fig. S14A**). LENT silencing was further associated with activation of the DNA damage response with increased gH2AX seen both by immunofluorescence (**Fig. S14B**) and immunoblot (**Fig. S14C**). Immunoblot analyses showed close to maximal LC3-II accumulation already 16 hours after LENT silencing, whereas gH2AX appeared only after 24 hours suggesting it was a secondary effect of mitophagy (**Fig. S14D**).

The observed autophagy/mitophagy and impaired OxPhos prompted us to investigate changes in translation of mitochondrial proteins of the electron transport complexes. Strikingly, increased protein levels of ATP5A, UQCR2, SDHB, COX II and NDUFB8 were seen after ASO-mediated LENT silencing in 501Mel cells and in melanocytic Mel888 and MM117 cells (**Fig. 8A and Fig S15)**. In contrast, this accumulation was not seen in LENOX silenced cells despite the fact that its silencing was also associated with lowered OxPhos capacity (15) (**Fig. 8A)**. Both mitophagy and OxPhos protein accumulation were therefore specific to LENT-silenced cells. Increased ATP5A levels, the most strongly affected in cells, was also seen in extracts from LENT ASO-treated IGR37 xenograft tumours (**Fig. 8B**).

**Figure 8:**
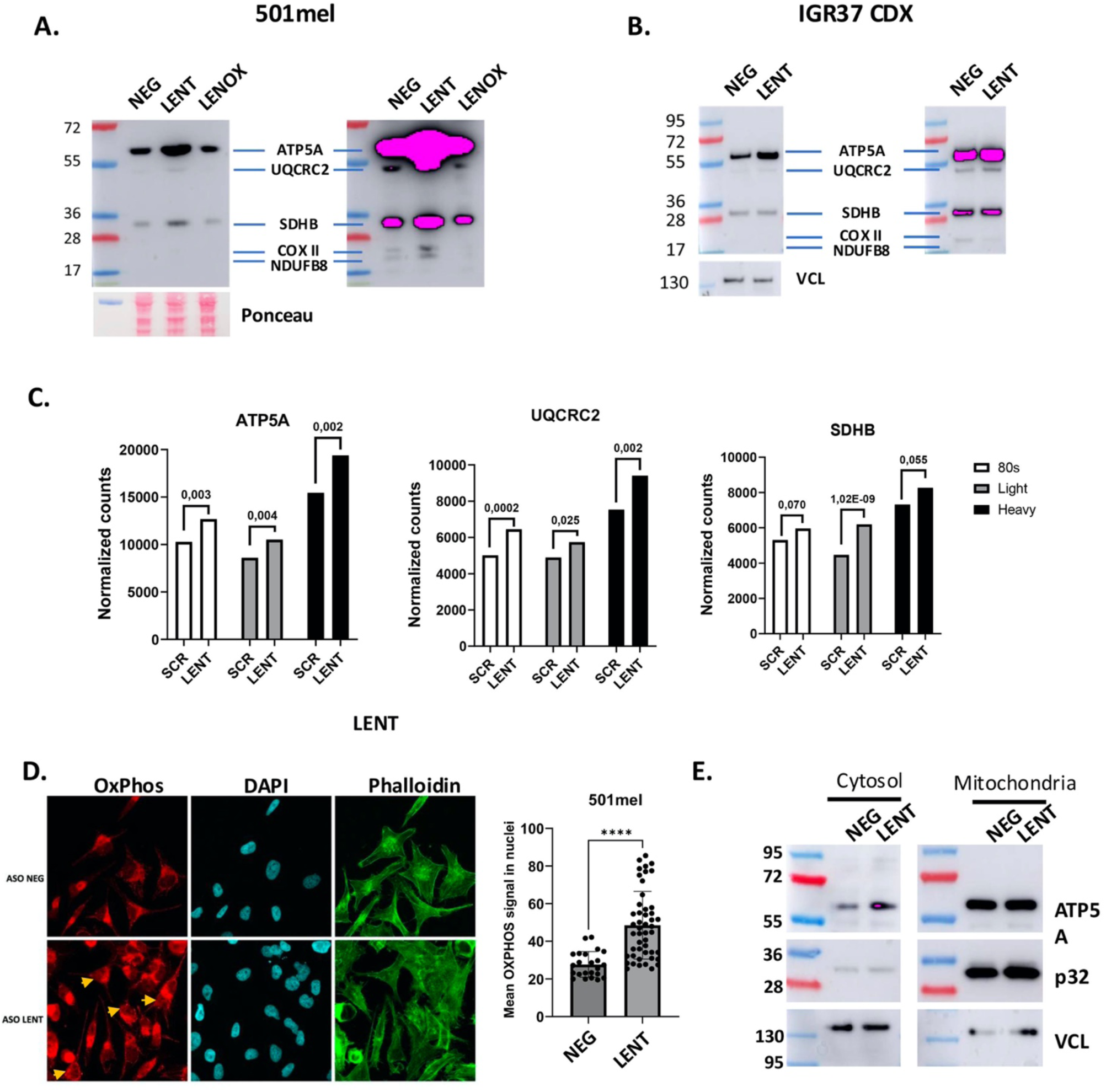
LENT depletion triggers DNA damage and accumulation of OxPhos proteins. **A.** Immunoblots detecting the indicated proteins in control or LENT depleted cells. Panels show different exposures of the same immuoblot. **B.** Immunoblots detecting mitochondrial electron transport chain proteins in extracts from IGR-37 CDX tumours. **C.** RNA-seq data showing the representation of the indicated RNAs in the 80S, LP and HP fractions. The normalized number of reads are shown along with the adjusted p-value between the control and shLENT conditions. **D.** OxPhos proteins were detected by confocal microscopy immunofluorescence in control or LENT-silenced cells using phalloidin to stain the cytoplasm. Proteins were quantified in the nucleus using the DAPI signal and comparisons were made by Mann-Whitney test. Scale bars = 10 µM. Representative nuclei displaying strong OxPhos protein staining are indicated by arrows. **E.** Immunoblot of ATP5A1 in cytosolic or mitochondrial fractions upon LENT depletion. Ponceau is shown as loading control. *, P < 0.033; **, P < 0.0021; ***, P < 0.0002; ****, P < 0.0001.

OxPhos protein accumulation may represent a compensatory response to the mitophagy and impaired mitochondrial function and/or may at least in part result from their increased translation as the presence of the corresponding mRNAs was up-regulated in the polysome fractions from shLENT silenced cells (**Fig. 8C**). However, as the increase in association with the LP and HP fractions was modest, their accumulation may also result from their impaired ERAD-mediated degradation. Furthermore, their accumulation was surprising given the lowered OxPhos, suggesting the excess OxPhos proteins were not imported into the mitochondria, but accumulated in the cytoplasm. Immunofluorescence showed accumulation of mitochondrial proteins at the mitochondria, but also in the cytoplasm and the nucleus of the LENT-silenced cells (**Fig. 8D**). Immunoblots on the cytosolic and mitochondrial fractions showed increased presence of ATP5A in the cytosolic fraction, with little change in the mitochondrial fraction showing that the accumulated protein was not imported into the mitochondria but rather accumulated outside the mitochondrial (**Fig. 8E**).

Together these data are consistent with the idea that LENT silencing impaired ER-homeostasis resulting in auto/mitophagy with subsequent impaired mitochondrial function. Accumulation of the mitochondrial proteins around the mitochondria and in the nucleus may then induce a general proteotoxic stress leading to apoptosis (30, 31, 32).

## Discussion

### LENT, a multi-functional lncRNA

Here we characterize LENT as a cytoplasmic lncRNA that interacts with the G4 resolvase DHX36 to promote translation of mRNAs involved in ER homeostasis and mitochondrial function and supressing autophagy in melanoma cells. LENT was primarily expressed in melanocytic, but not mesenchymal type melanoma cells. While a murine orthologue of LENT has been reported (37), we did not observe its expression in murine melanoma cells (data not shown). LENT expression did not correlate with poor survival in primary melanoma, but correlated with poor survival in metastatic melanoma where it was expressed in proliferative melanocytic type cells marked by an OxPhos signature (15, 33, 34). ASO-mediated LENT silencing induced apoptosis in cultured melanoma cells and impaired xenograft tumour growth. Moreover, simultaneous ASO targeting of LENT together with LENOX or SAMMSON cooperatively impacted melanoma cell viability. Thus ASO-targeting of these lncRNAs individually or in combination highlights their potential as therapeutic targets.

LINC00520 has been the focus of previous studies designated as LASSIE (35) or LEENE (36) LASSIE was described as a lncRNA induced in endothelial cells by sheer-stress that interacts with PECAM-1 to regulate vascular homeostasis by stabilizing adherens junctions. On the other hand, Miao et al (36) reported that LEENE was induced by pulsatile or oscillatory sheer stress in endothelial cells, but was localized in the nucleus and acted as an enhancer (e)RNA to regulate eNOS expression. In these non-melanoma cells, LINC00520 expression is driven by KLF2 and KLF4 highlighting that transcription factors other than MITF can drive its expression. Moreover, LEENE was further shown to promote transcription of pro-angiogenic genes, angiogenesis and tissue repair following ischemia (37). In contrast in melanoma cells, RNA-scope and cell fractionation showed that LENT was distributed between the cytosol and the mitochondria. We did not see enrichment of PECAM-1 in the RNA pulldown/mass-spectrometry experiments and LENT silencing did not affect NOS3 (eNOS) expression. This comparison between our data and that previously reported shows that LINC00520 is a multifunctional lncRNA (38) functioning in a cell-type and context dependent manner as an eRNA in the nucleus to regulate gene expression or in the cytoplasm to regulate translation.

### LENT coordinates ribosomal association of mRNAs encoding proteins involved in ER and mitochondrial homeostasis in melanocytic melanoma cells

We found that LENT selectively and directly interacted with the DHX36 G4 resolvase suggesting that it may contain a G4 structure. The *in vitro* interaction between LENT and DHX36 was reduced, but not abolished by mutation of a G-rich sequence with predicted potential to form a G4 structure. Nevertheless, this G-rich sequence diverges from more canonical G4-forming sequences and LENT does not comprise the 5’-GGnGGnGG-3’ motif or other motifs previously shown to be enriched in DHX36-associated RNAs (22, 27). G4 forming sequences are however variable with the length and sequence of the loop regions between the G blocks contributing to selectivity (39, 40) with DHX36 showing high specificity for parallel G4 structures (18). Moreover, LENT specifically pulled down DHX36, but no other well characterized G4 resolvases such as the RECQL-family including BLM and WRN, nor DDX5, DDX11, DHX9 or DDX3X (39, 40) (41) (24). While several of these helicases are mainly nuclear and more specific for G4 structures in DNA, DDX3X, DDX5 and DHX9 for example are reported RNA G4 resolvases (24, 41, 42). The selectivity of the LENT-DHX36 interaction may therefore reflect specific features of a potential LENT G4 sequence or alternatively, this interaction may be mediated by sequences or structures in LENT independent of G4-formation.

We provide evidence that LENT modulates DHX36 interaction with a subset of mRNAs. DHX36-RIP from control 501Mel cells identified DHX36-associated RNAs of which around 40% were previously identified as harboring G4 structures by other methods in other cell types. The most strongly associated RNAs were enriched in potential G4 forming motifs and the 5’-GGnGGnGG-3’ motif (27) (22). Surprisingly however, the overlap between the DHX36-associated RNAs in the HEK293T cells used by Sauer et al (27) and the melanoma cells was much lower than seen with the other G4 enrichment protocols, perhaps reflecting the different gene expression profiles in these cell lines. DHX36 RIP upon LENT silencing identified RNAs whose association with DHX36 was either increased or decreased. Increased binding may be explained if LENT were to comprise a G4 structure that simply competes with the G4s in other RNAs for DHX36 binding. However, this competition mechanism cannot explain the reduced binding of RNAs with DHX36 seen upon LENT silencing. How LENT binding to DHX36 modifies its interaction with these RNAs in a positive or negative manner remains to be determined.

An important observation of this study is the association of DHX36 with ribosomes, in the 80S and the polysome fractions. This observation contrasts with that of Sauer et al, where ribosomal DHX36 was not readily seen in HEK293T cells (27), but rather is in accordance with Murat et al, (41) who found DHX36 associated with the 80S and polysome fractions in HeLa cells. The additional presence of LENT in the 80S and light polysome fractions suggested, either that it was itself translated (LENT comprises several short ORFs) and/or that it may modulate the selectivity of DHX36 to resolve G4 structures in target mRNAs promoting/inhibiting their translation. In accordance with this idea, LENT silencing resulted in a selective depletion of a set of mRNAs in the LP fractions. Association of many of these mRNAs with the 80S and HP fractions was also reduced, but to a lesser extent. Of these, 118 showed reduced interaction with DHX36 upon LENT silencing. In contrast, several mRNAs whose association with DHX36 was up-regulated upon LENT silencing were enriched in the 80S, LP and HP fractions. Immunoblots showed that mRNAs whose engagement with the LP fractions was promoted by LENT were less well translated upon its silencing and *vice versa*.

Together the above results support the idea that LENT, via interaction with DHX36, positively or negatively regulates engagement of mRNAs with polysomes and fine-tunes their subsequent translation. However, while there was a strongly significant overlap, not all transcripts showing reduced association with the LP fraction displayed reduced interaction with DHX36 upon LENT silencing. These mRNAs were however enriched in G4 sequences suggesting they nevertheless required DHX36-driven unwinding for engagement in the LP fraction. It is possible that their interactions with DHX36 were less stable or that DHX36 only acts during the elongation steps of their translation, one they are engaged with the ribosome. Alternatively, we cannot formally exclude the existence of alternative DHX36-independent mechanisms by which LENT regulates their polysome engagement and translation. We note that it was previously reported that DHX36 silencing often had only minor effects on translation of its associated mRNAs. Quantitative mass-spectrometry revealed that DHX36 silencing had only marginal effects on the translation of its associated mRNAs (27). Here the effects are even more restricted as only a subset of DHX36 associated mRNAs were regulated by LENT, some positively leading to increased translation and *vice versa*.

Despite the above caveats, the idea that the LENT-DHX36 axis regulates translation is in accordance with previous studies reporting that DHX36 unwinds G4 structures in mRNA to regulate their association with ribosomes and their translation in HEK293T and HeLa cells (27) (41). Similarly, DHX36 binds and regulates translation of the mRNA encoding GNAI1 and other mRNAs involved in skeletal muscle stem cell function (22). In these studies, silencing of DHX36 was shown to fine tune translation suggesting that when silenced DHX36 function can be carried out by one or several of the above-mentioned helicases. We rather showed how the selectivity of DHX36 was regulated by LENT. Indeed, the changes in mRNA engagement with polysomes seen upon LENT silencing were more pronounced than those reported for DHX36-associated mRNAs in HeLa and HEK293T cells upon DHX36 silencing. Thus, when present, DHX36 acts as a predominant mRNA G4 resolvase whose activity was modulated by LENT.

LENT is not the first lncRNA shown to affect DHX36 activity. Matsumura et al, (43) identified a cytoplasmic G4-containing lncRNA designed GSEC that binds and inhibits DHX36 promoting motility of colon cancer cells. Similarly, SMaRT is a lncRNA that binds the G4 of the MLX-ψ isoform preventing unwinding by DHX36 and repressing its translation in murine muscle differentiation (44). Nevertheless, while GSEC and SMaRT seem to act as molecular decoys to inhibit DHX36 function, LENT is unique in its ability to both positively and negatively impact DHX36 function in an RNA selective manner.

### LENT suppresses autophagy to promote melanoma cell survival

As described above, LENT promotes engagement of a collection of mRNAs with the LP fractions. Strikingly, these mRNAs are strongly enriched in multiple aspects of ER and protein homeostasis, encoding numerous subunits of several protein complexes or pathways, such as SEL1-SYNV1 required for ERAD, TAP1, TAP2 and CALR involved in HLA transport and antigen presentation, enzymes and machinery involved protein glycosylation or the SEC61-NOMO multi-pass complex. Reduced translation of these mRNAs would be expected to lead to accumulation of mis-folded and/or mis-localized proteins and ER stress. For example, down-regulation of WFS1 has previously been shown to lead to reduced OxPhos capacity, increased mitochondrial-lysosome contact and mitophagy similar to what was observed here (45) (46). Consequently, a major phenotype of LENT silencing is autophagy and mitophagy that rapidly appeared in LENT silenced cells associated with impaired OxPhos capacity.

A further consequence of impaired ER function and mitophagy is accumulation of mitochondrial proteins in the cytoplasm and nucleus either via their increased translation and/or impaired ER-degradation. We note that DHX36 was detected within the mitochondria and hence has the potential to regulate translation of both nuclear and mitochondrial encoded transcripts, although this will require further investigation. Mitochondrial protein accumulation led to proteotoxic stress and finally apoptosis that may further involve activation of the DNA damage response. Our data therefore support the idea that the major function of LENT is to fine-tune translation of these mRNAs and optimize ER/protein homeostasis, maintain OxPhos capacity, suppress autophagy/mitophagy and promote survival and proliferation of melanocytic melanoma cells.

Previous studies showed that lncRNA SAMMSON acted to coordinate cytoplasmic and mitochondrial translation in melanoma cells to antagonize mPOS-mediated apoptosis (14) (13). Here we show that LENT also regulates translation, but via a different mechanism, to antagonize proteotoxic stress and autophagy-mediated apoptosis. Fine-tuning of translation and protein homeostasis therefore seem to be critical lncRNA-regulated processes in proliferative melanoma cells with a common feature being optimization of mitochondrial function and OxPhos capacity (47).

Why melanocytic melanoma cells, as opposed to mesenchymal melanoma cells where LENT is not expressed, specifically require fine-tuning of translation and high OxPhos capacity is unclear. However, one likely possibility is that melanocytic melanoma cells, like normal melanocytes, often synthesize melanin, a process that involves production of reactive oxygen species rendering them particularly vulnerable to oxidative stress (48) (49). Optimization of mitochondrial and ER function may therefore be essential to antagonize oxidative stress and may therefore explain why proliferative melanoma cells exploit such diverse mechanisms to optimize mitochondrial homeostasis and OxPhos capacity and ensure cell viability.

## Materials and Methods

### Analysis of the TCGA-SKCM cohort

For analysis of TCGA-SKCM, raw-counts were retrieved and primary tumors were separated from distant metastasis samples. The raw-counts matrices were normalized by sequencing depth using DESeq2 size-factors and then gene-counts were divided by median transcript length. Consensus clustering was done in R using the ConsensusClusterPlus v3.17 package following standard procedure. In short, matrices were filtered to keep only coding genes based on their biotype annotation and the 5000 most variable genes were selected with the mad() function. The matrices were median centered with sweep(), apply() and median() functions before performing consensus clustering with ConsensusClusterPlus() using base parameters. The number of clusters were selected based on the curve of cumulative distribution function in order to define 4 clusters for primary tumors (CCP1-CCP4) and 5 clusters for distant metastasis samples (CCM1-CCM5).

### Cell culture and transfections

Melanoma cell lines SK-MEL-25, SK-MEL-25R (gifts from Dr L. Larue), SK-MEL-28, and 501mel (ATCC) were grown in RPMI1640 w/o HEPES medium supplemented with 10 % fetal calf serum (FCS) and gentamycin (40 µg/mL); IGR-37 and IGR-39 in RPMI1640 w/o HEPES medium supplemented with 15% FCS and gentamycin (40 µg/mL). MM011, MM117, MM047, and MM099 (gifts from Dr J-C. Marine) were grown in HAM-F10 medium supplemented with 10 % FCS, 5.2 mM glutamax, 25 mM Hepes, and penicillin/streptomycin (7.5 μg/mL). M229, M229R, M249, and M249R (gifts from Dr J-C. Marine) were grown in DMEM medium supplemented with glucose (4.5 g/L), 5 % FCS, and penicillin/streptomycin (7.5 μg/mL). A375 cells were grown in DMEM medium supplemented with glucose (4.5 g/L), 10 % FCS, and gentamycin (40 µg/mL). HEK293T cells were grown in DMEM medium supplemented with glucose (1 g/L), 10 % FCS, and penicillin/streptomycin (7.5 ug/mL). HeLa cells were grown in DMEM medium with glucose (1 g/L), 5 % FCS and gentamycin (40 µg/mL). To assess cell growth and viability, cells were stained with Trypan Blue (Invitrogen). Trametinib (GSK1120212) and dabrafenib (GSK2118436) were purchased from Selleckchem, Salubrinal was purchased from Merck. SK-MEL-25, Sk-MEL-28, A375, and 501mel were obtained from ATCC, all other cell lines were gifts from collaborators. All cell lines were regularly tested using the Venor GeM Mycoplasma Detection Kit, and used at less than 10 passages. ASO and siRNA were transfected using Lipofectamine RNAiMAX (Invitrogen) with 20 nM of ASO (Qiagen) or siRNA (Thermo Fisher Scientific). ASO and siRNAs sequences are listed below. For ASO combination experiments, cells were transfected with 15 nM of LENT ASO and/or 15 nM of LENOX ASO and/or 5 nM of SAMMSON ASO. For trametinib+dabrafenib-GapmeR and salubrinal-GapmeR cotreatment, cells were cultured for 3 days in presence or absence of Dabrafenib (100 nM) + Trametinib (100 nM) or Salubrinal (20 µM), transfected with 15 nM of GapmeR and then cultured for additional 3 days before harvesting. Colony-forming ability was assessed by plating 500 cells/9.6 cm2, wait for 10 days, fixing cells in formalin and staining with 0.05 % Crystal Violet solution (Sigma Aldrich).

### CRISPR interference

501mel cells were co-transfected with a plasmid expressing dead Cas9 protein fused to the Kruppel-associated box (KRAB) domain-containing KAP1 (dCas9-KAP1) and the red fluorescent protein mScarlet (pX-dCas9-KRAB-Scarlet), together with another plasmid expressing GFP and three single guide RNAs targeting the transcription start site of LENT (pcDNA3-sgRNA-GFP) or a control plasmid expressing GFP only (pCMV-GFP). Double Scarlet-GFP positive cells were sorted 24 hours after co-transfection, stained with Cell Trace Violet and cultured for additional 96 hours.

### Plasmid cloning and lentiviral transduction

For the ectopic expression experiment, LENT cDNA was cloned into the pCW57-GFP-P2A-MCS vector (a gift from Adam Karpf; Addgene plasmid #71783; http://n2t.net/addgene:71783; RRID: Addgene_71783). LENT shRNA (shLENT) or a scrambled control (shCTRL) were cloned in LT3GEPIR (a gift from Johannes Zuber; Addgene plasmid #111177; http://n2t.net/addgene:111177; RRID: Addgene_111177). Lentiviral particles were produced in HEK293T cells, purified by ultracentrifugation, and resuspended in PBS. Lentiviruses were titrated with flow cytometry by measuring the GFP signal intensity in HEK293T infected with different dilutions of viruses. Melanoma cells were eventually infected at a multiplicity of infection (MOI) of 1 and selected by puromycin addition to the media (1 mg/mL) in every following passage.

### RNAscope

LENT and MITF RNAs were detected with the RNAscope assay (Advanced Cell Diagnostics, ACD) according to the manufacturer’s protocol. Patient sections were deparaffinized, incubated with hydrogen peroxide at room temperature for 10 minutes, boiled with target retrieval reagent for 15 minutes, and then treated with protease plus reagent at 40 °C for 30 minutes. Sections were hybridized with Hs-MITF probe (ACD, catalog no. 310951) and hs-LENT at 40 °C for 2 hours. Probes for Hs-LENT were custom designed by ACD. Hybridization signals were amplified and visualized with RNAscope Multiplex Fluorescent Reagent Kit v2 (ACD, catalog no. 323100). For co-detection of DHX36 with LENT, cells were fixed for 30 minutes with formaldehyde 3.7 %, washed with PBS and incubated 10 minutes at room temperature with H2O2. After one wash in distilled water, primary antibody for DHX36 diluted in co-detection diluent (1/200) was added o/n at 4 °C. Slides were washed in PBS + tween 0.1 % (PBST), fixed in formaldehyde 3.7 % for 30 minutes, and washed again in PBST. Slides were treated with protease III and washed with PBS. LENT hybridization signals were amplified following the Multiplex Fluorescent Kit. Finally, DHX36 signal was developed by secondary antibody incubation (diluted 1/2,000 in co-detection diluent), followed by tyramide signal amplification (TSA Plus Kit, NEL760001KT, Perkin Elmer). Images were captured with a confocal (Leica DMI6000) microscope. Mander’s and Pearson’s coefficients were calculated with the Fiji software using the JACoP plugin.

### Analysis of oxygen consumption rate in living cells

Oxygen consumption rate (OCR) was measured in an XF96 extracellular analyzer (Seahorse Bioscience). 20,000 transfected cells per well were seeded 48 hours prior the experiment. The cells were incubated at 37°C and the medium was changed to XF base medium supplemented with 1 mM pyruvate, 2 mM glutamine, and 10 mM glucose for 1 hour before OCR profiling with the Mitostress Test Kit sequentially exposed to 2 μM oligomycin, 1 μM carbonyl cyanide-p-trifluoromethoxyphenylhydrazone (FCCP), and 0.5 μM rotenone and antimycin A. Cells were washed with PBS, fixed with 3 % PFA and permeabilized with 0.2 % triton. Nuclei were counterstained with DAPI (1:500) and number of cells per well was determined with a Celomics Cell Insight CX7 (Thermofisher Scientific). Non-mitochondrial oxygen consumption was subtracted from the total values to designate true mitochondrial oxygen consumption values.

### Flow cytometry

To assess cell viability and proliferation, cells were stained with Cell Trace Violet (Invitrogen) on the day of transfection, harvested after 72 hours and stained with Annexin V (BioLegend) and TOPRO-3 (Invitrogen) or the active caspase-3 Kit (BD Biosciences). Cell Trace Violet staining was performed following the manufacturer’s instructions and as previously described (15)(50). All analyses were performed on a LSRII Fortessa (BD Biosciences) and data were analyzed with FlowJo software (TreeStar).

To analyse intracellular ROS, cells were stained in adherent conditions with CellRox Deep Red (Thermo Fisher Scientific) at final concentration of 500 nM following manufacturer instructions. After harvesting, cells were stained for active caspase-3 (BD Biosciences) and analyzed on a LSRII Fortessa (BD Biosciences). To induce reactive oxygen species (ROS), cells were treated with THBP (200 μM) for 30 minutes. To induce apoptosis, cells were treated with staurosporine (500 nM) for 16 hours.

### LENT pulldown and LC/MS-MS analysis

501mel cells were grown in 15 cm petri dishes, harvested by trypsinization, washed, pelleted, resuspended in lysis buffer (TrisHCl 20 mM pH8, NaCl 200 mM, MgCl2 2.5 mM, Triton 0.05%, DEPC water) supplemented with fresh DTT (1 mM), protease and phosphatase inhibitor cocktail (Thermo Fisher Scientific) and RNAsin (Thermo Fisher Scientific) and kept 20 minutes on ice. For crosslinked pulldown, petri dishes were exposed to 400 mJ/cm2 of UV radiation with a CL-1000 crosslinker (254 nm lamp) and the concentration of NaCl in lysis buffer was adjusted to 300 mM. Membranes were pelleted at 3,000 g for 3 minutes at 4°C and supernatant precleared for 1 hour at 4 °C with 100 µg of streptavidin-coated sepharose beads (Cytiva). The lysate was incubated 2 hours with streptavidin coated beads and 400 pmol anti-PCA3 or LENT-specific DNA biotinylated oligonucleotides (listed below). Beads were pelleted for 3 minutes at 3,000 g and washed five times with lysis buffer. After final wash beads were divided for RNA and protein extraction. RNA was purified by TRI Reagent and isopropanol precipitation, digested with DNAse, reverse transcribed and analyzed by qPCR. Proteins were eluted by boiling beads in Laemmli sample buffer and separated on NuPAGE Novex 4% to 12% gradient gels. For mass spectrometry analysis, three independent experiments were performed and the entire lane was excised after staining with Simply blue safe stain solution (Invitrogen). Analysis was performed at the Harvard Medical School Taplin Mass Spectrometry Facility, as described previously (9, 15).

### IGR37 xenograft model and ASO treatment

Swiss nude mice were purchased from Charles River Laboratories (France) and housed under specific pathogen-free conditions. Animal care, use, and experimental procedures were conducted in accordance with recommendations of the European Community (86/609/EEC), European Union (2010/63/UE) and the French National Committee (87/848). The ethics committee of IGBCM in compliance with institutional guidelines approved animal care and use (APAFIS#2023010611181767). Mice were injected on the rear flank with 3x106 IGR37 cells resuspended in 100 µL of 1x PBS + Cultrex Basement Membrane Extract (ref. 3432–005–01; R&D Systems) with a 1:1 ratio. Tumor growth was monitored by caliper measurement every two days and volume was calculated with the formula: (4/3 π) * (length/2) * (width/2) * (height/2). After tumors reached 100 mm3, mice were injected subcutaneously around 1 cm from the tumour every two days with 15 mg/kg of ASO for the LENT group, or not injected for the control group. After two weeks of treatment, mice were sacrificed and primary tumors were dissected and mechanically lysed in TRI Reagent for RNA extraction or LSDB for protein extraction.

### Immunofluorescence of fixed and live cells

Cells grown on Millicell EZ slides (Millipore) were fixed with 4 % paraformaldehyde for 15 minutes. After two washes with PBS buffer, they were permeabilized in PBS + Triton X-100 0.1 % for 5 minutes and blocked with PBS + 10 % FCS for 20 minutes. Primary antibodies were incubated overnight at 4 °C and after three washes with PBS + Triton 0.1%, cells were stained for 1 hour at room temperature with Alexa Fluor-488 conjugated secondary antibodies (Life technologies) diluted 1/500 in PBS + 10 % FCS. After three washes with PBS + Triton 0.1%, cells were stained with DAPI (final concentration 1 μg/mL) and mounted on microscopy slides with Prolong Gold antifade reagent (Invitrogen). Anti-DHX36 (13159-1-AP) and anti-HSP60 were diluted 1/200 in PBS + 10 % FCS. Images were captured with a confocal (Leica DMI6000) microscope. DHX36 enrichment at mitochondria was calculated with the ratio of the DHX36 signal overlapping with HSP60 signal over the total DHX36 signal for each cell on field.

For lysotracker + mitotracker experiment, live cells were incubated in medium complemented with Lysotracker Deep Red 1/20 000, Mitotracker Green FM 1/10 000 and Hoechst 33342 1/10 000 for 1 h, washed and then observed with a confocal microscope inside a chamber at 37 °C with 5 % CO2.

### Transmission Electron Microscopy and immune-gold staining

Samples were fixed by immersion in 2.5 % glutaraldehyde and 2.5 % paraformaldehyde in cacodylate buffer (0.1 M, pH 7.4), washed in cacodylate buffer for further 30 minutes. The samples were postfixed in 1% osmium tetroxide in 0.1 M cacodylate buffer for 1 hour at 4 °C and dehydrated through graded alcohol (50, 70, 90, and 100%) and propylene oxide for 30 minutes each. Samples were oriented and embedded in Epon 812. Semithin sections were cut at 2 µm and ultrathin sections were cut at 70 nm (Leica Ultracut UCT) and contrasted with uranyl acetate and lead citrate and examined at 70 kv with a Morgagni 268D electron microscope (FEI Electron Optics, Eindhoven, the Netherlands). Images were captured digitally by Mega View III camera (Soft Imaging System). Immuno-gold labeling was performed using AURION immunogold reagents essentially as described: (https://aurion.nl/labeling-protocol/post-embedding/protocol-conventional-reagents/) <u>using the DHX36 and COX IV antibodies</u>.

### RNA extraction and RT-qPCR

Total RNA isolation was performed using TRI Reagent (MRC) and isopropanol precipitation, according to the manufacturer protocol. Pelleted RNAs were resuspended in water and DNA was depleted using the TurboDnase Free Kit (Thermo Fisher Scientific). RNA was then reverse transcribed with the Superscript IV reverse transcriptase (Thermo Fisher Scientific) following manufacturer instructions. qPCR was carried out with SYBR Green I (Roche) and monitored by a LightCycler 480 (Roche). Target gene expression was normalized using TBP, HBMS, and RPL13A as reference genes. For polysome profiling normalization was performed using mRNAs encoding GAPDH, TBP and HMBS. Primers for RT-qPCR are listed below.

### Protein extraction and Western blotting

Whole cell extracts were prepared by freeze–thaw technique using LSDB 500 buffer [500 mM KCl, 25 mM Tris at pH 7.9, 10 % glycerol (v/v), 0.05 % NP-40 (v/v), 16 mL DTT, and protease inhibitor cocktail]. Lysates were subjected to SDS-PAGE and proteins were transferred onto a nitrocellulose membrane. Membranes were incubated with primary antibodies in PBS + 5 % BSA + 0.01 % Tween-20 o/n at 4°C. The membrane was then incubated with HRP-conjugated secondary antibody (Jackson ImmunoResearch, 1/2000) for 1 hour at room temperature, and visualized using the ECL detection system (GE Healthcare). Antibodies used are listed below.

### Mitochondria fractionation

Mitochondria were isolated with the Mitochondria Isolation Kit (Thermo Fisher Scientific) following manufacturer instructions. Briefly, harvested cells were washed and pelleted, resuspended in buffer A, and incubated 2 minutes on ice. Buffer B was added for 5 minutes, vortexing every minute, and diluted with buffer C. Nuclei were pelleted 10 minutes at 700 × g and supernatant centrifuged for 15 minutes at 3,000 × g. Purified mitochondria were washed in buffer C and lysed in CHAPS 2%.

For mitochondrial protein content analysis after digitonin treatment, mitochondria were purified as described above. Then, purified mitochondria were digested on ice for 15 minutes with increasing concentrations of Digitonin (Invitrogen) in mitochondria isolation buffer (210 mM Mannitol, 70 mM Sucrose, 1 mM EDTA, 10 mM HEPES and protease inhibitors cocktail). Digested mitochondria were pelleted by centrifugation (13 000 g for 10 minutes) and the supernatant was removed. Digested mitochondria were then lysed and the protein content was analyzed by SDS-PAGE.

For the trypsin treatment, mitochondria were also prepared following the Mitochondria Isolation kit. Mitochondria were subsequently digested 20 minutes on ice with 50 µg/mL Trypsin diluted in mitochondria isolation buffer. Digestion was stopped by adding 120 µg / mL of soybean trypsin inhibitor. Pellets were centrifuged 10 minutes at 13 000 g at 4 °C, and lysed as described above. For cell swelling conditions, mitochondria were digested with trypsin diluted in HEPES-KOH 20 mM at pH 7.5 instead of mitochondria isolation buffer.

### Identification of RNAs associated with DHX36

Cells were grown in 15 cm petri dishes, harvested by scraping, resuspended in lysis buffer (20 mM Tris-HCl pH 8, 200 mM NaCl, 2.5 mM MgCl2, 0.05 % Triton, DEPC water) supplemented with DTT (1 mM), protease/phosphatase inhibitor cocktail (Thermo Fisher Scientific) and RNAsin (Thermo Fisher Scientific) and kept on ice for 15 minutes, pipetting every 3 minutes. Membranes were pelleted 10 minutes at 10,000 g at 4 °C and the supernatant precleared 1 hour at 4 °C with protein G magnetic beads (Invitrogen). Lysate was quantified by Bradford protein quantification assay (Bio-Rad) and incubated overnight on a rotating wheel at 4 °C with 5 µg of the indicated antibodies. Then, 50 µL of resuspended Protein G magnetic beads were added for 3 hours to isolate RNA – protein complexes and washed five times in lysis buffer. After final wash, RNA was purified by TRI Reagent + isopropanol precipitation and proteins eluted by boiling beads at 95 °C for 15 minutes in Laemmli buffer.

For RNA sequencing, RNAs were obtained from 501mel cells expressing a control shRNA or an shRNA targeting LENT following the method described above. RNA profiles were determined by using a 2100 Bioanalyser. rRNAs were depleted with the Ribo-Zero Plus rRNA depletion kit (Illumina) and the libraries were prepared with the library prep mRNA ultralow Smarter kit (Takara). Sequencing was performed on a NextSeq 2000 high throughput sequencer (Illumina). Analyses were performed as described in the next paragraph.

### Bulk RNA sequencing data

Gene expression in 501mel cells transfected with control or LENT-targeting ASO2 was analyzed by RNA-seq. After sequencing raw reads were pre-processed in order to remove adapter and low-quality sequences (Phred quality score below 20) using cutadapt version 1.10. and reads shorter than 40 bases were discarded. Reads were mapping to rRNA sequences using bowtie version 2.2.8, were also removed. Reads were mapped onto the hg19 assembly of Homo sapiens genome using STAR version 2.5.3a. Gene expression quantification was performed from uniquely aligned reads using htseq-count version 0.6.1p1, with annotations from Ensembl version 75 and “union” mode. Only non-ambiguously assigned reads were retained for further analyses. Read counts were normalized across samples with the median-of-ratios method. Comparisons of interest were performed using the Wald test for differential expression and implemented in the Bioconductor package DESeq2 version 1.16.1. Genes with high Cook’s distance were filtered out and independent filtering based on the mean of normalized counts was performed. P-values were adjusted for multiple testing using the Benjamini and Hochberg method.

### Motif enrichment analysis

To identify RIP-seq genes containing G-quadruplex regions, we first retrieved publicly available RIP-seq data for DHX36 reported by Varshney et al (51) from GEO (accession: GSE154570). We then performed de-novo motif analysis using the MEME-ChIP algorithm on DHX36 RIP retained RNAs identifying the G-quadruplex motif “CCGCCGCY” and generating the associated probability matrix. Lastly, we used the FIMO algorithm to find this motif in the 5’UTR regions of genes identified in the DHX36 RIP-seq analysis.

### RNA in-vitro transcription and purification

Double-stranded DNA molecules (gBlocks) containing LENT WT or LENT ΔG sequences were ordered from Integrated DNA Technologies and used as PCR templates to generate RNA by in-vitro transcription. DNA constructs were designed with the T7 RNA polymerase promoter sequence. After run-off transcription, RNAs were purified by denaturing polyacrylamide gel electrophoresis (PAGE) and extracted by the “crush and soak method” as described (52). For EMSA experiments, the purified RNA transcripts were labelled at their 5ʹ end by addition of a radioactive cap using the Vaccine Capping Enzyme with the ScriptCap m7G capping system from CELLSCRIPT in the presence of [32P] αGTP (>6000 Ci/mmol). The 5ʹ-radiolabelled transcript was separated from enzyme and free nucleotides by Bio-Spin 6 Columns (Biorad).

### DHX36 Protein Expression and Purification

The inducible expression plasmid (pDHX36 54-989), encompassing the sequence of the human protein (aa 54 to 898) with a His6-SUMO N-terminus tag (53), was generously provided by Rick Russell. Recombinant protein expression was carried out in BL-21 DE3 Rosetta2 pLysS cells (Merck). Cells were grown at 37 °C until reaching an OD-600 nm value of 0.9. Subsequently, the temperature was lowered to 18 °C, and protein expression was induced by adding 0.5 mM IPTG for 16 hours. Cells were harvested by centrifugation, and the pellet was resuspended in lysis buffer [50 mM Tris-HCl pH 8.0, 1 M NaCl, 10 % Glycerol, 10 mM 2-Mercaptoethanol, 1 mM CaCl2, 10 µg/mL Dnase I (Merck), 1x Halt-Protease (Pierce)] before being lysed by sonication at 4 °C. The lysate was clarified by centrifugation, and nucleic acids were precipitated using 0.1 % polyethyleneimine and removed by centrifugation. The supernatant was then passed through a Ni-NTA column (Protino, Macherey-Nagel) equilibrated with buffer A (50 mM Tris-HCl pH 8.0, 1 M NaCl, 10 % Glycerol). After extensive washing with the equilibration buffer, the protein was eluted with buffer A supplemented with 300 mM Imidazole. Factions containing the protein were treated with ULP Protease at a ratio of 1:500 (W/W) and digested/dialyzed overnight at 4 °C in buffer B (50 mM Tris-HCl pH 8.0, 350 mM NaCl, 10 mM 2-Mercaptoethanol, 10 % Glycerol). The DHX36 54-989 protein, liberated from its N-terminal tag, was separated on a NiNTA column equilibrated with buffer B, with the protein predominantly found in the flow-through fractions. Further purification was accomplished via chromatography on a Heparin affinity column (Hitrap HP, Cytiva). The DHX36 54-989 protein was eluted by a linear gradient from 10 to 500 mM NaCl in buffer (50 mM Tris-HCl pH 8.0, 10 % Glycerol, 10 mM 2-Mercaptoethanol), resulting in a symmetrical peak, and isolated at the end of the gradient. The protein fractions were concentrated in the elution buffer to a final concentration of 7 µM (concentration determined using OD at 280 nm, extinction coefficient 108180 M-1cm-1), snap-frozen in liquid nitrogen, and stored at -80°C. The identity of the final purified protein was confirmed by mass spectrometry analysis (performed at Strasbourg-Esplanade Proteomics Facility). Dynamic light scattering (DLS) was used to investigate the solubility of the purified protein.

### EMSA

5’ end labeled LENT WT and/or LENT ΔG RNA (10,000 cpm; < 3 nM) and a molar excess of oligo(dT) in 8 μl of milli-Q (Millipore) water were heated for 2 min at 90 °C and chilled on ice for 2 min. After addition of tenfold concentrated refolding buffer (50 mM MES/NaOH pH 6, 100 mM KOAc, 1 mM Mg(Oac)2 and one unit of Rnasin (Promega), RNA was renatured for 15 min at RT. 10 µl of RNA was finally incubated 30min in ice with increasing concentrations of DHX36 (0-125nM) in twofold concentrated binding buffer (final concentration: 50mM Tris/HCl pH8, 150mM NaCl, 10% glycerol, 10mM β-mercaptoethanol). Electrophoresis was performed in TBM (89 mM Tris base, 89 mM boric acid and 1 mM Mg(Oac)2) buffer at 120 V for 5 h at 4 °C. Results were analyzed by phosphorimaging. Quantitative analysis was performed using ImageLab software (Biorad).

### Sucrose density gradient ribosome profiling

Cells were washed twice with PBS, harvested by scraping and rapidly centrifuged at 300 g for 5 min at 10 °C. The resulting pellet was resuspended in lysis buffer (20 mM HEPES/NaOH pH 7.4, 100 mM KOAc, 1 mM DTT, 0.5 mM Mg(Oac)2, 100 U of Recombinant Rnasin (Promega) and Halt Protease inhibitor cocktail (ThermoFisher)). Cell lysis was performed by nitrogen cavitation with 4639 Cell Disruption Vessel (Parr Instrument Company) at 350 psi for 50 min, stirring with a small magnet at 500 rpm in a cold room (4 °C). Lysate was then centrifuged at 1000 g for 5 minutes and the supernatant was recovered avoiding the foam (membranes) and the pellet (nuclei). After an incubation of 5 min at 30 °C, 20–30 OD260 of cell extracts from both cell lines were gently layered over 7–47 % sucrose gradients in buffer T (25 mM Hepes/NaOH pH 7.4, 79 mM KOAc, 2.5 mM Mg(Oac)2, 1 mM DTT, 3 U/µl Rnasin (Promega). Gradients were centrifuged at 37000 rpm (Beckman, SW41Ti) for 2 h and 30 min at 4 °C. After centrifugation, 45 fractions (0.25 ml/fraction) were collected on a BIOCOMP gradient fractionator equipped with an UV detector. For RNA-seq, biological triplicate ribosome profiling was performed from control or shLENT-silenced cells as described above. RNA from three fractions corresponding to the 80S, light or heavy polysome fractions was pooled and sequenced using the Ribo-Zero Plus rRNA depletion kit (Illumina) as described above.

### Statistics

All tests used for statistical significance were calculated using GraphPad Prism10 and indicated in the figure legends along with p values (****p < 0.0001, ***p < 0.001, **p < 0.01, *p < 0.05, ns: p > 0.05).

## Resources

All oligonucleotides and antibodies used are listed in Tables 1-4 below.

## Data availability

The datasets generated during and/or analyzed in this study are available from the corresponding author I. Davidson upon request. The RNA-seq data described in this paper have been deposited with the GEO data base under the accession number GSE270716.

### ASO and shRNA

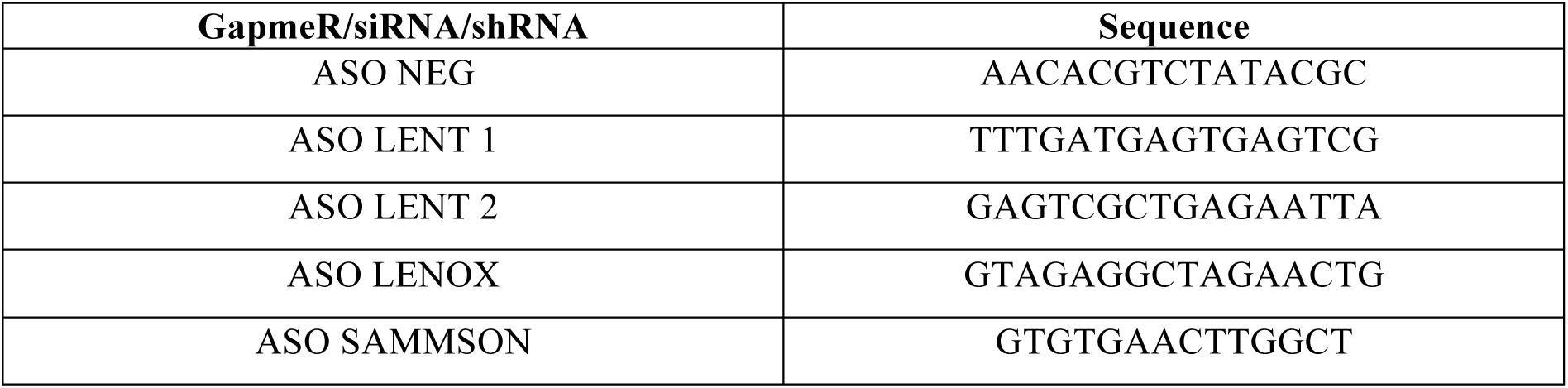
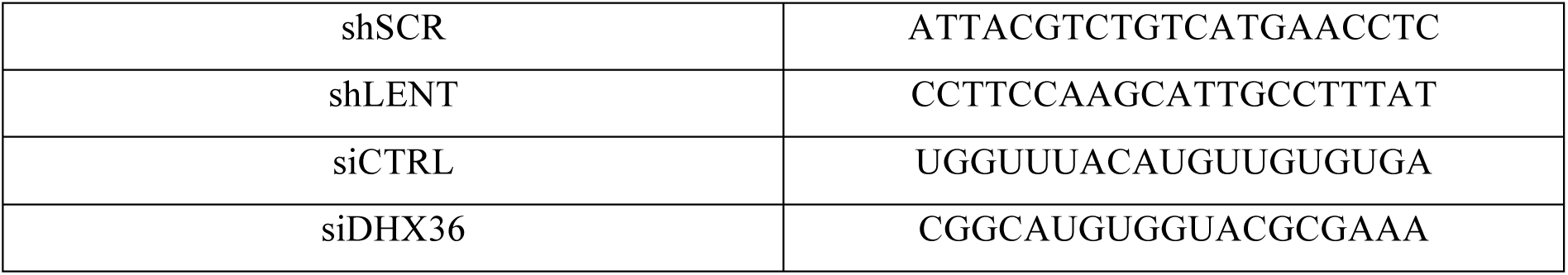

### Primers

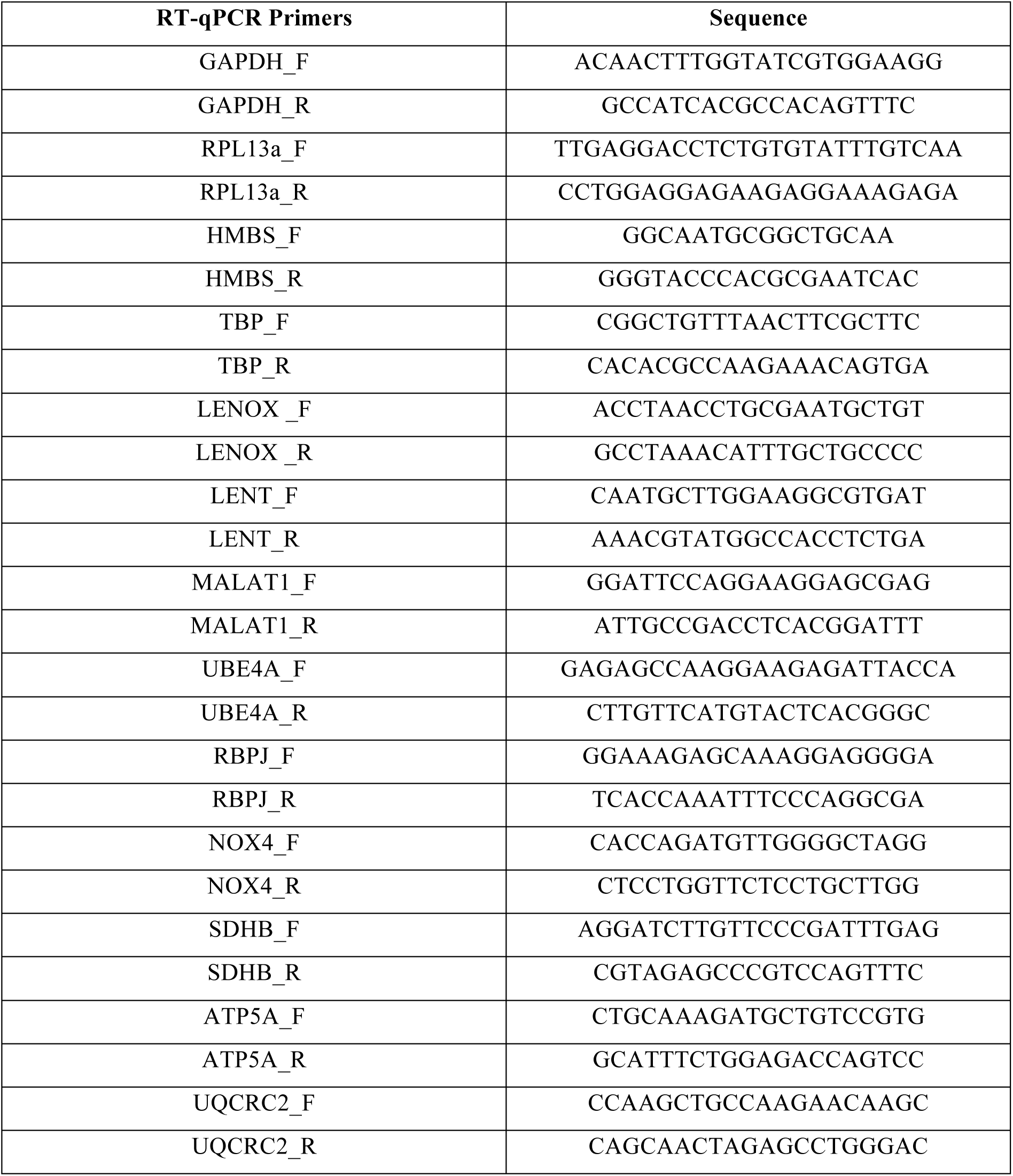

### Antibodies

| Primary Antibodies | Host | Application | Dilution | Lot number |
| --- | --- | --- | --- | --- |
| DHX36 (13159-1-AP) | Rabbit | WB | 1/1000 | 00004147 |
|  |  | IF<br>IP | 1/200<br>5 µg/mL |  |
| CARF (BE-A303-861A-M) | Rabbit | WB | 1/1000 | 1 |
| RAP2 (sc-515711) | Mouse | WB | 1/1000 | B4397 |
| HSP60 (in house) | Mouse | WB<br>IF | 1/500<br>1/100 | 4MTE- 2H7 |
| VINCULIN (V4505) | Mouse | WB | 1/5000 | 099M4850V |
| ACTIN (in house) | Mouse | WB | 1/1000 | 1ACT- 2D7 |
| COX IV (ab202554) | Rabbit | WB | 1/1000 | GR3342068-1 |
| UBE4A (sc-365904) | Mouse | WB<br>IF | 1/1000<br>1/200 | C3117 |
| NOX4 (NB110-58849) | Rabbit | WB<br>IF | 1/1000<br>1/200 | D134519 |
| EIF4A2 (ab31218) | Rabbit | WB | 1/1000 | GR3383097-1 |
| RPL36 (PA5-117106) | Rabbit | WB | 1/2000 | WG3327290E |
| C1QBP (A302-863A) = p32 | Rabbit | WB<br>IF | 1/1000<br>1/200 | 1 |
| TOMM20 (H00009804-M01) | Mouse | WB | 1/1000 | HC5202153D |
| RBP-Jk (sc-271128) | Mouse | WB<br>IF | 1/500<br>1/100 | A2220 |
| OXPHOS (45-8199) | Mouse | WB | 1/1000 | VB2939036 |
| LC3B (ab51520) | Rabbit | WB | 1/5000 | GR3374012-2 |
| gH2AX (ab22551) | Mouse | WB<br>IF | 1/1000<br>1/100 | GR3358071-2 |
| FUNDC1 (NBP1-81063) | Rabbit | WB | 1/500 | 000042870 |
| CTSD (2284) | Rabbit | WB | 1/1000 | 2 |
| WFS1 (26995-1-AP) | Rabbit | WB | 1/500 | 00096359 |
| HSPA5 (HPA038845) | Rabbit | WB | 1/500 | A83196 |
| Normal Rabbit IgG (12-370) | Rabbit | IP | 5 µg/mL | 3493998 |

### Biotinylated oligonucleotides

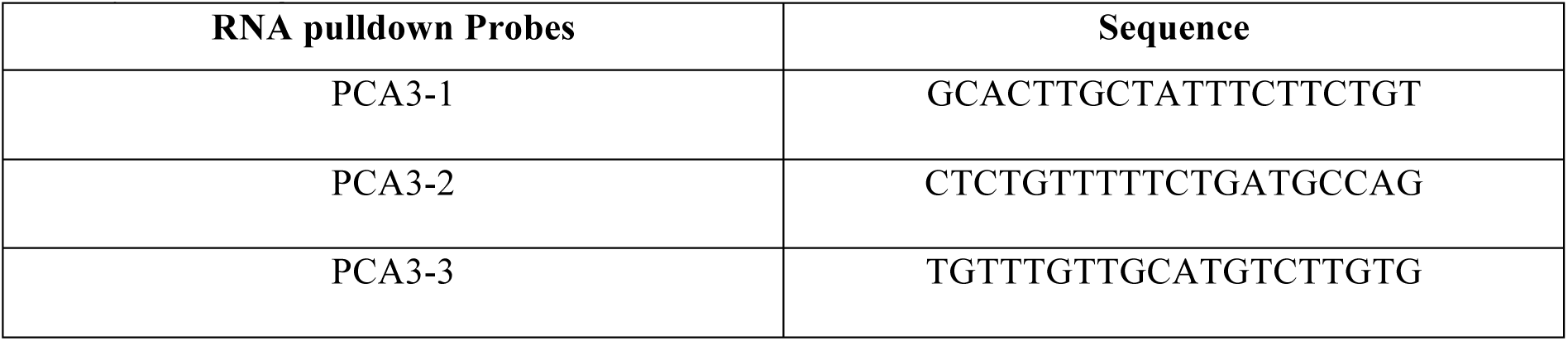
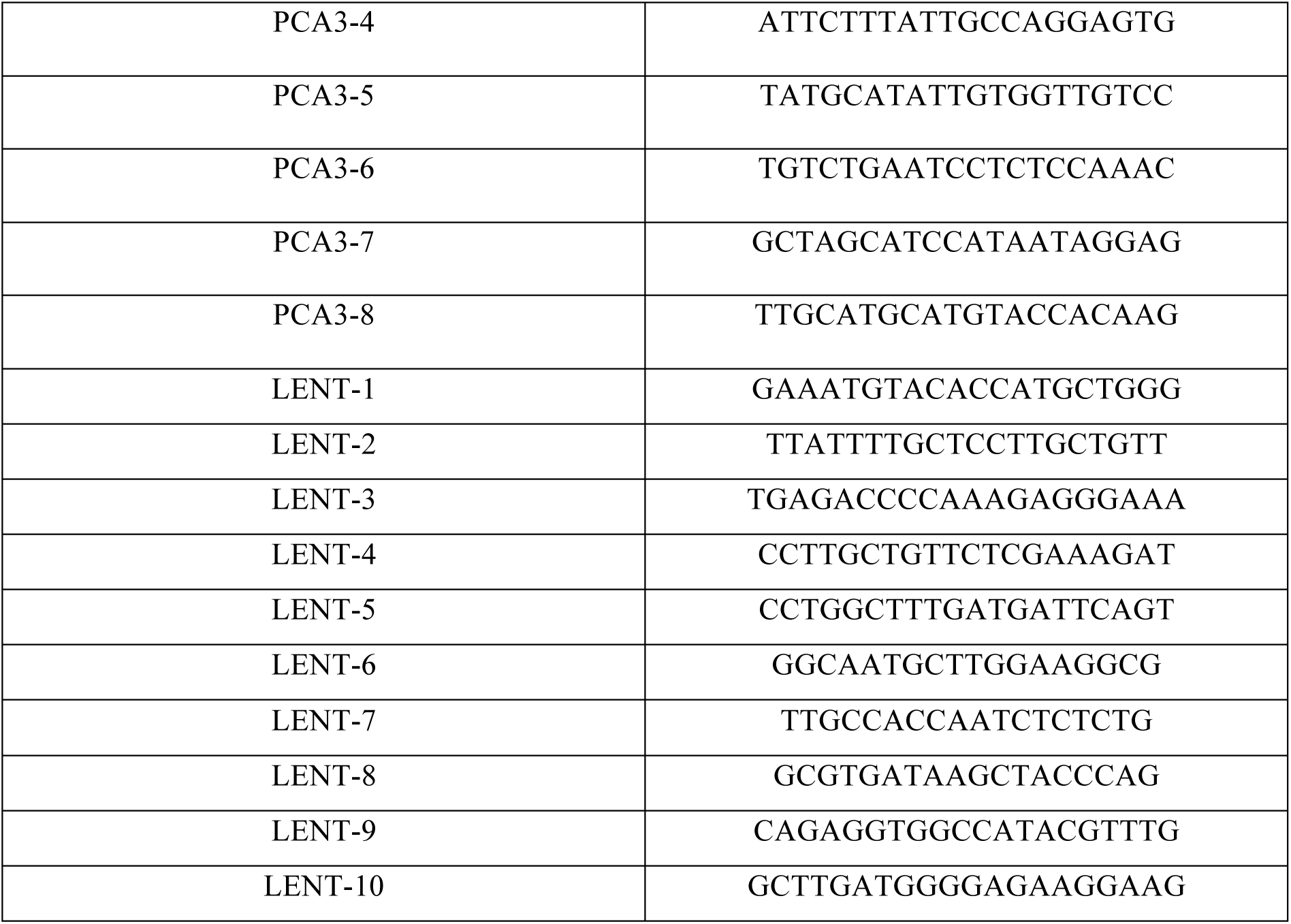

## Acknowledgements

We thank, I. Michel for excellent technical assistance, R Tomaino and the Harvard mass-spectrometry platform, L. Larue, J-C. Marine and G. Ghanem for the melanoma lines and MM-series primary melanoma cells, D. Lipkser and the Dermatology Clinic of Strasbourg University Hospital for patient melanoma sections, the GenomEast high throughput sequencing platform, the staff of the IGBMC common facilities. This work was supported by grants from the ITMO Cancer, the Ligue Nationale contre le Cancer, SATT Conectus Alsace, the ANR-10-LABX-0030 and ANR-10-IDEX-0002-02. ID is an ‘équipe labellisée’ of the Ligue Nationale contre le Cancer. AH was supported by a fellowship from the Fondation pour la Recherche Médicale.

## Author Contributions

AH, GG and GM performed all the ASO targeting cell culture and xenograft experiments, GD, AH and GG performed bioinformatics analyses, AH performed the LENT RNA pulldown and subsequent functional experiments, Md’A, GB, AS and EE generated in vitro transcribed LENT, recombinant DHX36, performed the EMSA and the polysome profiling, CN and NM performed the electron microscopy and immune-gold staining, AH, GG, EE and ID conceived the experiments, analysed the data and wrote the paper.

## References

1. Tirosh I, et al. Dissecting the multicellular ecosystem of metastatic melanoma by single-cell RNA-seq. Science. 2016;352(6282):189–96.

2. Tsoi J, et al. Multi-stage Differentiation Defines Melanoma Subtypes with Differential Vulnerability to Drug-Induced Iron-Dependent Oxidative Stress. Cancer Cell. 2018;33(5):890–904 e5.

3. Rambow F, et al. Toward Minimal Residual Disease-Directed Therapy in Melanoma. Cell. 2018;174(4):843–855 e19.

4. Rambow F, Marine JC, Goding CR. Melanoma plasticity and phenotypic diversity: therapeutic barriers and opportunities. Genes Dev. 2019;33(19–20):1295–1318.

5. Ennen M, et al. MITF-High and MITF-Low Cells and a Novel Subpopulation Expressing Genes of Both Cell States Contribute to Intra- and Intertumoral Heterogeneity of Primary Melanoma. Clin Cancer Res. 2017;23(22):7097–7107.

6. Karras P, et al. A cellular hierarchy in melanoma uncouples growth and metastasis. Nature. 2022;610(7930):190–198.

7. Pozniak J, et al. A TCF4-dependent gene regulatory network confers resistance to immunotherapy in melanoma. Cell. 2024;187(1):166–183.e25.

8. Strub T, et al. Essential role of microphthalmia transcription factor for DNA replication, mitosis and genomic stability in melanoma. Oncogene. 2011;30(20):2319–32.

9. Laurette P, et al. Transcription factor MITF and remodeller BRG1 define chromatin organisation at regulatory elements in melanoma cells. Elife. 2015;10.7554/eLife.06857. 10.7554/eLife.06857.

10. Verfaillie A, et al. Decoding the regulatory landscape of melanoma reveals TEADS as regulators of the invasive cell state. Nat Commun. 2015;6:6683.

11. Mauduit D, et al. Analysis of long and short enhancers in melanoma cell states. Elife. 2021;10:e71735.

12. Marin-Bejar O, et al. Evolutionary predictability of genetic versus nongenetic resistance to anticancer drugs in melanoma. Cancer Cell. 2021;39(8):1135–1149.e8.

13. Vendramin R, et al. SAMMSON fosters cancer cell fitness by concertedly enhancing mitochondrial and cytosolic translation. Nat Struct Mol Biol. 2018;25(11):1035–1046.

14. Leucci E, et al. Melanoma addiction to the long non-coding RNA SAMMSON. Nature. 2016;531(7595):518–22.

15. Gambi G, et al. The lncRNA LENOX interacts with RAP2C to regulate metabolism and promote resistance to MAPK inhibition in melanoma. Cancer Res. 2022;CAN-22-0959.

16. Trotta AP, et al. Disruption of mitochondrial electron transport chain function potentiates the pro-apoptotic effects of MAPK inhibition. J Biol Chem. 2017;292(28):11727–11739.

17. Luan W, et al. Long non-coding RNA LINC00520 promotes the proliferation and metastasis of malignant melanoma by inducing the miR-125b-5p/EIF5A2 axis. J Exp Clin Cancer Res. 2020;39(1):96.

18. Chen MC, et al. Structural basis of G-quadruplex unfolding by the DEAH/RHA helicase DHX36. Nature. 2018;558(7710):465–469.

19. Kikin O, D’Antonio L, Bagga PS. QGRS Mapper: a web-based server for predicting G-quadruplexes in nucleotide sequences. Nucleic Acids Res. 2006;34(Web Server issue):W676–682.

20. Antcliff A, McCullough LD, Tsvetkov AS. G-Quadruplexes and the DNA/RNA helicase DHX36 in health, disease, and aging. Aging. 2021;13(23):25578–25587.

21. Chen X, et al. Translational control by DHX36 binding to 5’UTR G-quadruplex is essential for muscle stem-cell regenerative functions. Nat Commun. 2021;12(1):5043.

22. Chen X, et al. Translational control by DHX36 binding to 5’UTR G-quadruplex is essential for muscle stem-cell regenerative functions. Nat Commun. 2021;12(1):5043.

23. Chen X, et al. Lockd promotes myoblast proliferation and muscle regeneration via binding with DHX36 to facilitate 5’ UTR rG4 unwinding and Anp32e translation. Cell Rep. 2022;39(10):110927.

24. Herdy B, et al. Analysis of NRAS RNA G-quadruplex binding proteins reveals DDX3X as a novel interactor of cellular G-quadruplex containing transcripts. Nucleic Acids Research. 2018;46(21):11592–11604.

25. Guo JU, Bartel DP. RNA G-quadruplexes are globally unfolded in eukaryotic cells and depleted in bacteria. Science. 2016;353(6306):aaf5371.

26. Kwok CK, et al. rG4-seq reveals widespread formation of G-quadruplex structures in the human transcriptome. Nat Methods. 2016;13(10):841–844.

27. Sauer M, et al. DHX36 prevents the accumulation of translationally inactive mRNAs with G4-structures in untranslated regions. Nat Commun. 2019;10(1):2421.

28. Govindarajan B, et al. Overexpression of Akt converts radial growth melanoma to vertical growth melanoma. J Clin Invest. 2007;117(3):719–729.

29. Arnoult D, et al. An N-terminal addressing sequence targets NLRX1 to the mitochondrial matrix. Journal of Cell Science. 2009;122(17):3161–3168.

30. Wang X, Chen XJ. A cytosolic network suppressing mitochondria-mediated proteostatic stress and cell death. Nature. 2015;524(7566):481–484.

31. Coyne LP, Chen XJ. MPOS is a novel mitochondrial trigger of cell death – implications for neurodegeneration. FEBS Letters. 2018;592(5):759–775.

32. Karbowski M, Oshima Y, Verhoeven N. Mitochondrial proteotoxicity: implications and ubiquitin-dependent quality control mechanisms. Cell Mol Life Sci. 2022;79(11):574.

33. Haq R, et al. Oncogenic BRAF regulates oxidative metabolism via PGC1alpha and MITF. Cancer Cell. 2013;23(3):302–15.

34. Vazquez F, et al. PGC1alpha expression defines a subset of human melanoma tumors with increased mitochondrial capacity and resistance to oxidative stress. Cancer Cell. 2013;23(3):287– 301.

35. Stanicek L, et al. Long non-coding RNA LASSIE regulates shear stress sensing and endothelial barrier function. Commun Biol. 2020;3(1):265.

36. Miao Y, et al. Enhancer-associated long non-coding RNA LEENE regulates endothelial nitric oxide synthase and endothelial function. Nat Commun. 2018;9(1):292.

37. Tang X, et al. Long noncoding RNA LEENE promotes angiogenesis and ischemic recovery in diabetes models. Journal of Clinical Investigation. 2023;133(3):e161759.

38. Mattick JS, et al. Long non-coding RNAs: definitions, functions, challenges and recommendations. Nat Rev Mol Cell Biol. 2023;24(6):430–447.

39. Varshney D, et al. The regulation and functions of DNA and RNA G-quadruplexes. Nat Rev Mol Cell Biol. 2020;21(8):459–474.

40. Lejault P, et al. How to untie G-quadruplex knots and why? Cell Chemical Biology. 2021;28(4):436–455.

41. Murat P, et al. RNA G-quadruplexes at upstream open reading frames cause DHX36- and DHX9-dependent translation of human mRNAs. Genome Biol. 2018;19(1):229.

42. Wu G, et al. DDX5 helicase resolves G-quadruplex and is involved in *MYC* gene transcriptional activation. Proc Natl Acad Sci USA. 2019;116(41):20453–20461.

43. Matsumura K, et al. The novel G-quadruplex-containing long non-coding RNA GSEC antagonizes DHX36 and modulates colon cancer cell migration. Oncogene. 2017;36(9):1191–1199.

44. Martone J, et al. SMaRT lncRNA controls translation of a G-quadruplex-containing mRNA antagonizing the DHX36 helicase. EMBO Reports. 2020;21(6):e49942.

45. Cagalinec M, et al. Role of Mitochondrial Dynamics in Neuronal Development: Mechanism for Wolfram Syndrome. PLoS Biol. 2016;14(7):e1002511.

46. Patergnani S, et al. The Wolfram-like variant WFS1E864K destabilizes MAM and compromises autophagy and mitophagy in human and mice. Autophagy. 2024;1–12.

47. Huang C, Radi RH, Arbiser JL. Mitochondrial Metabolism in Melanoma. Cells. 2021;10(11):3197.

48. Obrador E, et al. Oxidative stress and antioxidants in the pathophysiology of malignant melanoma. Biol Chem. 2019;400(5):589–612.

49. Denat L, et al. Melanocytes as Instigators and Victims of Oxidative Stress. Journal of Investigative Dermatology. 2014;134(6):1512–1518.

50. Coassolo S, et al. Citrullination of pyruvate kinase M2 by PADI1 and PADI3 regulates glycolysis and cancer cell proliferation. Nat Commun. 2021;12(1):1718.

51. Varshney D, et al. RNA G-quadruplex structures control ribosomal protein production. Sci Rep. 2021;11(1):22735.

52. Petrov A, et al. RNA Purification by Preparative Polyacrylamide Gel Electrophoresis. Methods in Enzymology. Elsevier; 2013:315–330.

53. Yangyuoru PM, et al. The G-quadruplex (G4) resolvase DHX36 efficiently and specifically disrupts DNA G4s via a translocation-based helicase mechanism. Journal of Biological Chemistry. 2018;293(6):1924–1932.

