## Supplemental Figures and Legends for "Interaction of lncRNA LENT with DHX36 regulates translation and suppresses autophagy in melanoma"

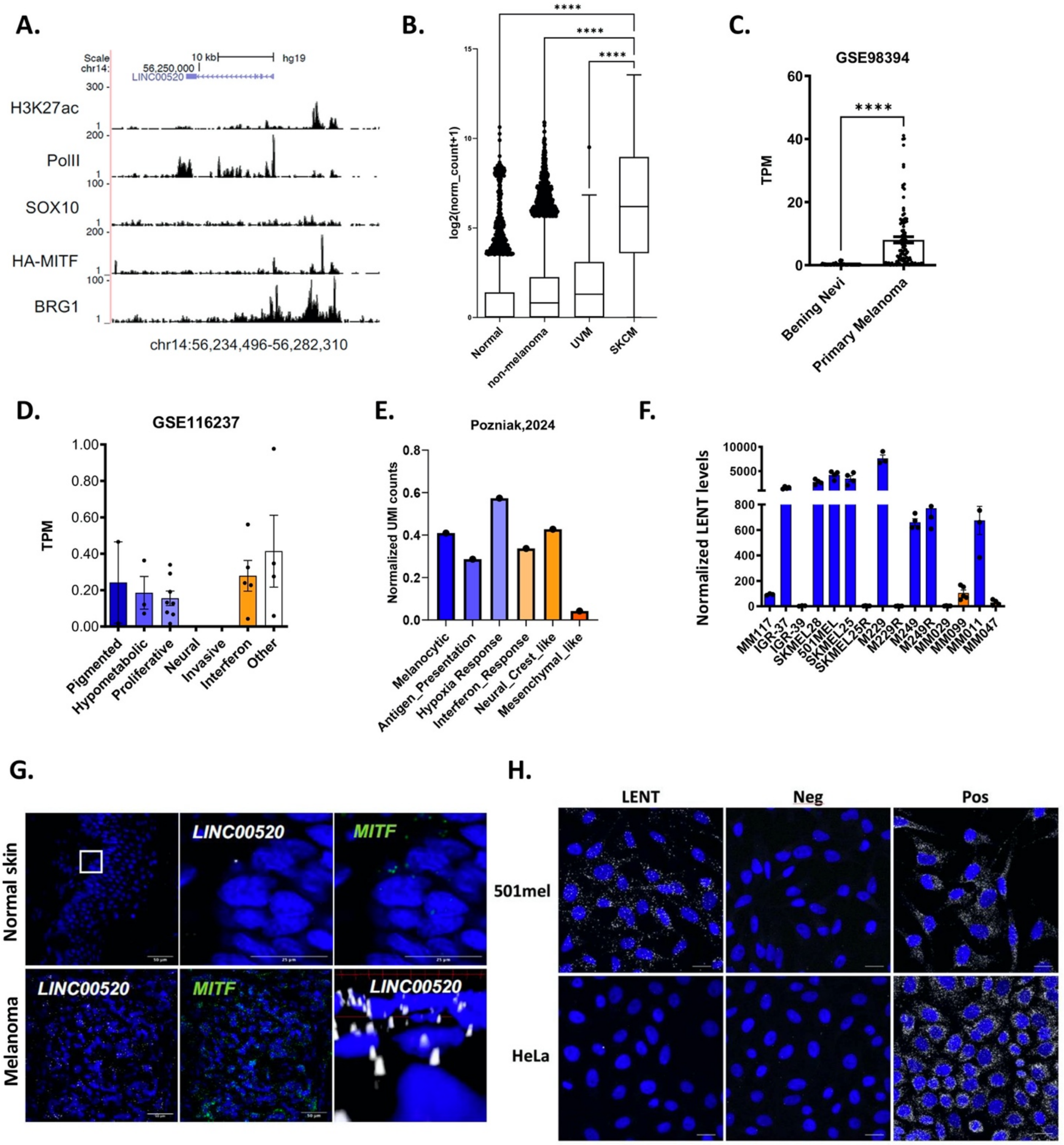
A.
Scale
chr14:
10 kb
hg19
56,250,000
LINC00520
H3K27ac
PolII
SOX10
HA-MITF
BRG1
chr14:56,234,496-56,282,310
B.
log2(norm_count+1)
Normal
non-melanoma
UVM
SKCM
****
C.
GSE98394
TPM
****
Bening Nevi
Primary Melanoma
D.
GSE116237
TPM
Pigmented
Hypometabolic
Proliferative
Neural
Invasive
Interferon
Other
E.
Pozniak,2024
Normalized UMI counts
Melanocytic
Antigen_Presentation
Hypoxia Response
Interferon_Response
Neural_Crest_like
Mesenchymal_like
F.
Normalized LENT levels
MM117
IGR-37
IGR-39
SKMEL28
501MEL
SKMEL25
SKMEL25R
M229
M229R
M249
M249R
MM029
MM099
MM011
MM047
G.
Normal skin
Melanoma
LINC00520
MITF
H.
LENT
Neg
Pos
501mel
HeLa

### Figure S1

**Figure S1: LENT expression is the highest in melanocytic cells of skin cutaneous melanoma. A.** UCSC screenshot of the indicated ChIP-seq tracks from Laurette et al (9) at the LENT locus illustrating binding of MITF and BRG1. **B.** LENT expression in tumors and normal tissues were retrieved from the GTEX and TCGA databases and compared by one-way ANOVA (Dunnett test). GTEX; N = 858. TCGA Non-melanoma: N = 9986; Uveal melanoma; N = 79. SKCM N = 470. **C.** RNA-seq data from GSE98394 shows LENT levels in benign nevi compared to melanoma by Mann-Whitney test. Nevi, N = 54; Primary melanoma, N = 101. **D.** scRNA-seq data from GSE116237 shows LENT expression in the different cell populations. Melanocytic subtypes are shown in blue and mesenchymal subtypes in orange. **E.** scRNA-seq data from EGAD00001009291 from Pozniak et al (7) shows LENT expression in the different cell populations. **F.** LENT levels were determined by RT-qPCR in a collection of melanoma cell lines. Melanocytic are in blue and mesenchymal in orange. **G.** RNAscope coupled with confocal microscopy to detect LENT and MITF mRNA in sections from normal skin and melanoma. Nuclei are stained with DAPI. Scale bars are shown on each image. **H.** RNAscope coupled with confocal microscopy to detect LENT RNA in 501Mel or HeLa cells along with the negative and positive control reactions provided by the supplier. Nuclei are stained with DAPI. Comparisons were made by one-way ANOVA (Dunn's test). *, P < 0.033; **, P < 0.0021; ***, P < 0.0002; ****, P < 0.0001.

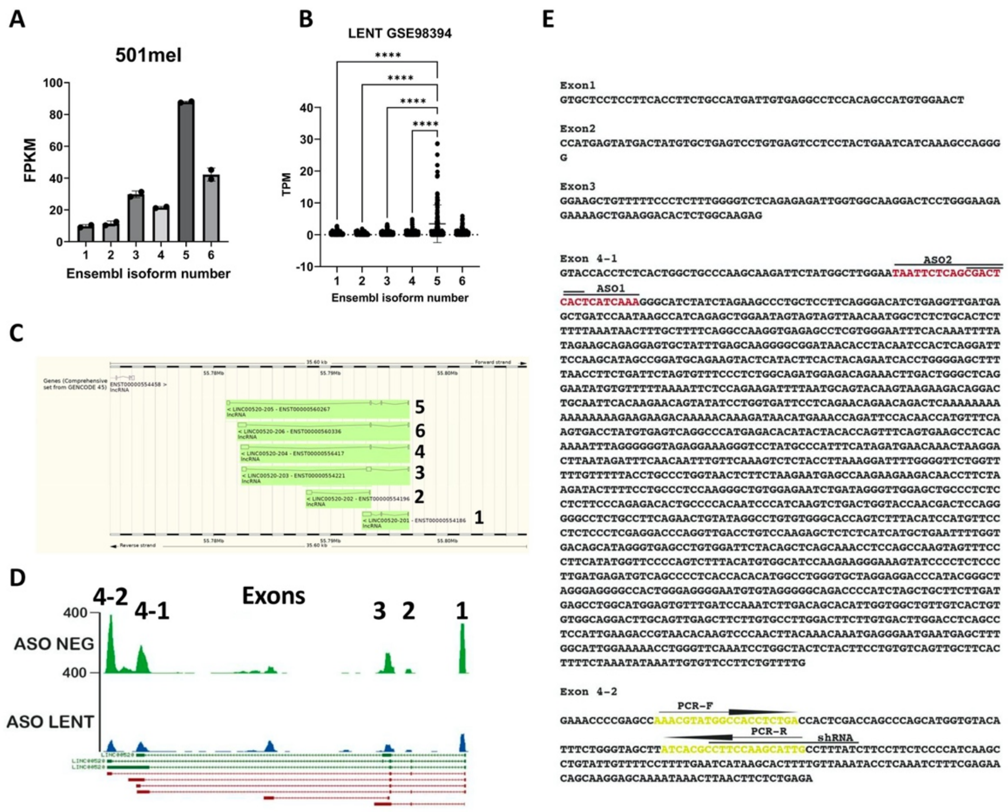


Figure S2

**Figure S2: LENT isoform expression. A-B.** LENT isoform expression in melanoma cell lines and in RNA-seq from patients (GSE98394). **C.** Isoforms are represented with the Ensembl genome browser interface. **D** RNA-seq track from 501Mel cells transfected with ASO control or LENT-targeting ASO2 illustrating exon coverage at the LENT locus. **E.** Sequences of the major exons of LENT. Exon 4-2 represents an internal splice acceptor in the last exon as illustrated in panel D. The sequences of the alternative exons of isoform 2 not found in the major isoforms are not shown.

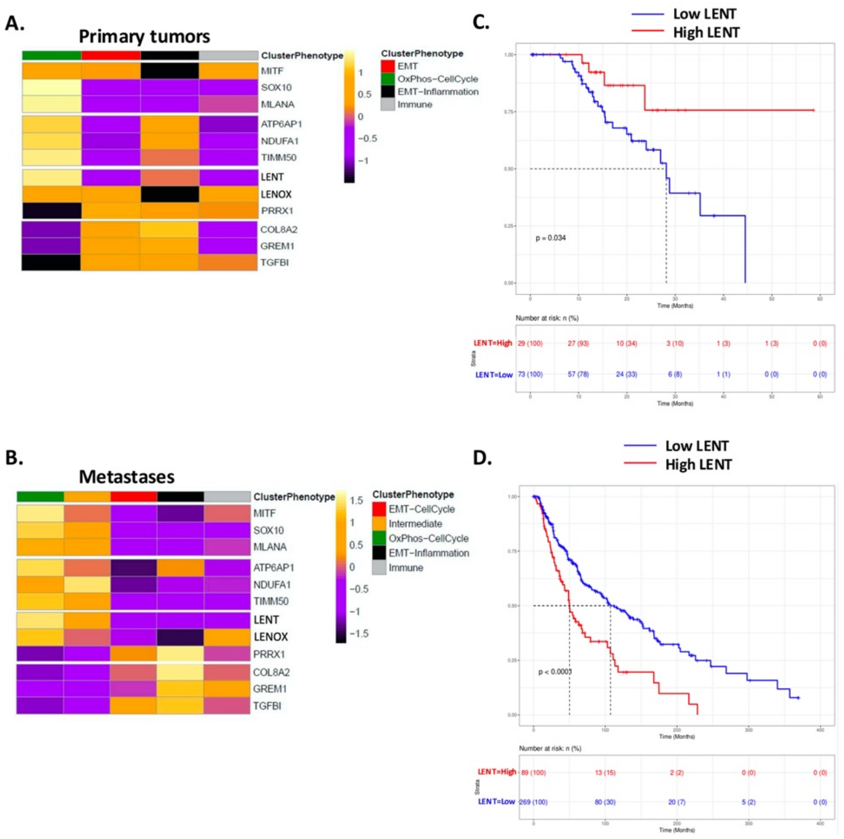


**Figure S3**

**Figure S3: LENT expression is highest in OxPhos enriched cells and correlates with melanoma patient survival. A and C.** Data from the SKCM TCGA were separated into primary tumor and metastases. Unsupervised clustering of the RNA-seq data from the two groups was performed and the identity of the cell populations derived from GSEA Hallmark analyses of the differentially expressed genes. Heatmaps show the expression of the indicated genes as a Z-scored heatmap in the identified cell populations. **B and D.** Kaplan-Meier curves for overall survival in patients according to LENT expression score using the optimal cut point method with the associated log-rank p-value from the univariate Cox proportional-hazard model. The number of patients in each group are indicated.

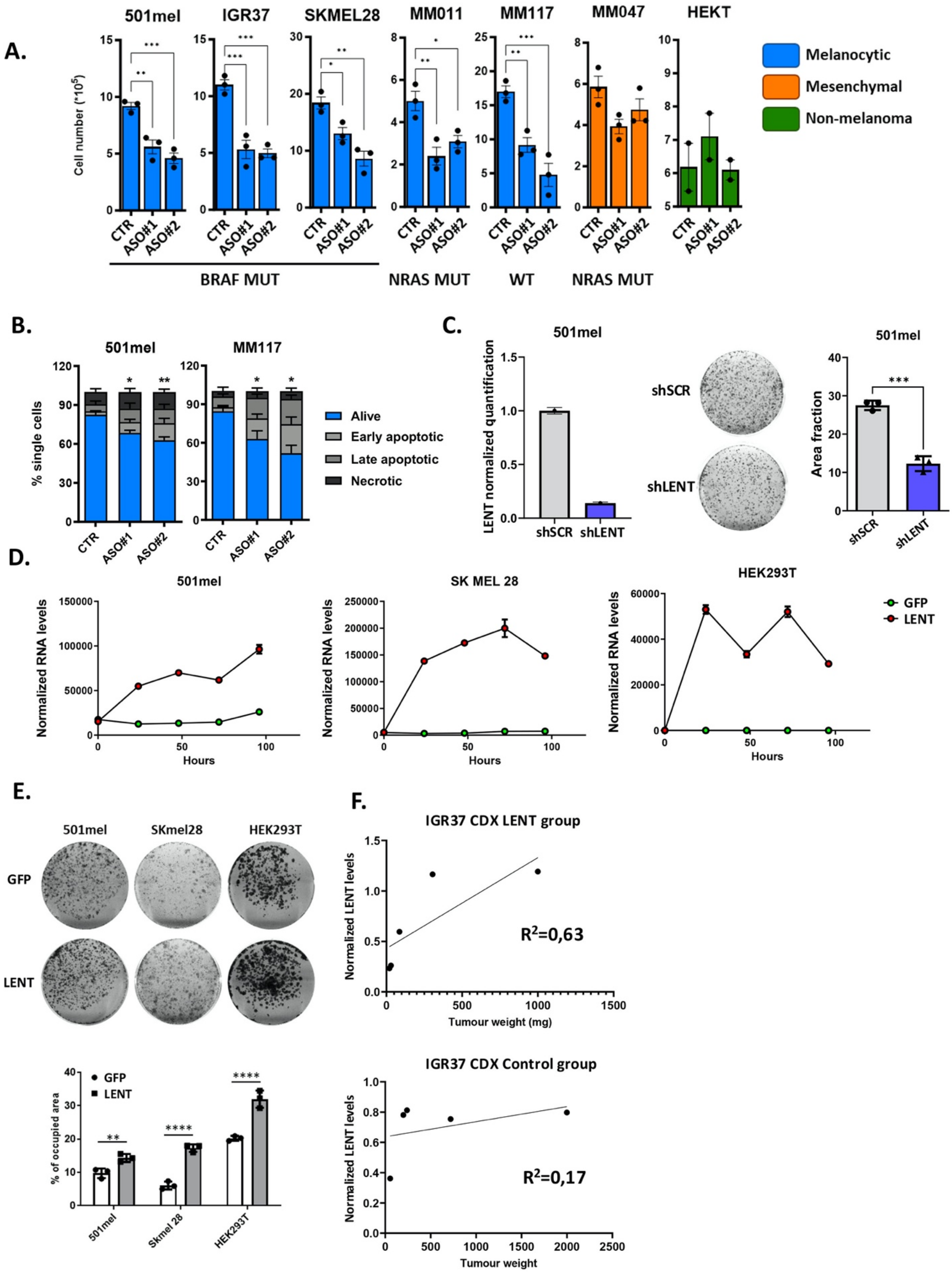
A.
501mel
IGR37
SKMEL28
MM011
MM117
MM047
HEKT
Cell number (*10^5)
CTR
ASO#1
ASO#2
BRAF MUT
NRAS MUT
WT
NRAS MUT
Melanocytic
Mesenchymal
Non-melanoma
B.
501mel
MM117
% single cells
Alive
Early apoptotic
Late apoptotic
Necrotic
C.
501mel
LENT normalized quantification
shSCR
shLENT
Area fraction
D.
501mel
SK MEL 28
HEK293T
Normalized RNA levels
Hours
GFP
LENT
E.
501mel
SKmel28
HEK293T
GFP
LENT
% of occupied area
Skmel 28
F.
IGR37 CDX LENT group
Normalized LENT levels
Tumour weight (mg)
R²=0,63
IGR37 CDX Control group
Tumour weight
R²=0,17

**Figure S4**

36

**Figure S4: Gain and loss of LENT in melanoma cells. A.** ASO-mediated depletion of LENT was performed in the indicated cell lines with two independent ASOs and cell numbers were counted after 48h and compared with a control condition by one-way ANOVA (Dunnett test). **B.** ASO-mediated depletion of LENT triggers apoptosis as shown by annexin V + PI staining coupled with flow cytometry and compared with a control condition by one-way ANOVA (Dunnett test). **C.** LENT was silenced by a Dox-inducible shRNA and LENT levels quantified by RT-qPCR. Comparison of colony formation following Dox-induced expression of control or LENT-targeting shRNA by unpaired T test. **D.** Ectopic expression of LENT in the indicated cell lines was measured by RT-qPCR using a cell line with GFP inducible as a control. **E.** Colony formation upon ectopic LENT expression was performed in the indicated cell lines and compared the GFP control by one-way ANOVA (Dunnett test). **F.** LENT levels in the IGR-37 CDX were measured by RT-qPCR and association with tumour weight was performed by linear regression. **G.** Cell proliferation was assessed after ASO-mediated LENT silencing in combination with the indicated drugs. Comparisons were done by two-way ANOVA. *, $P < 0.033$; **, $P < 0.0021$; ***, $P < 0.0002$; ****, $P < 0.0001$.

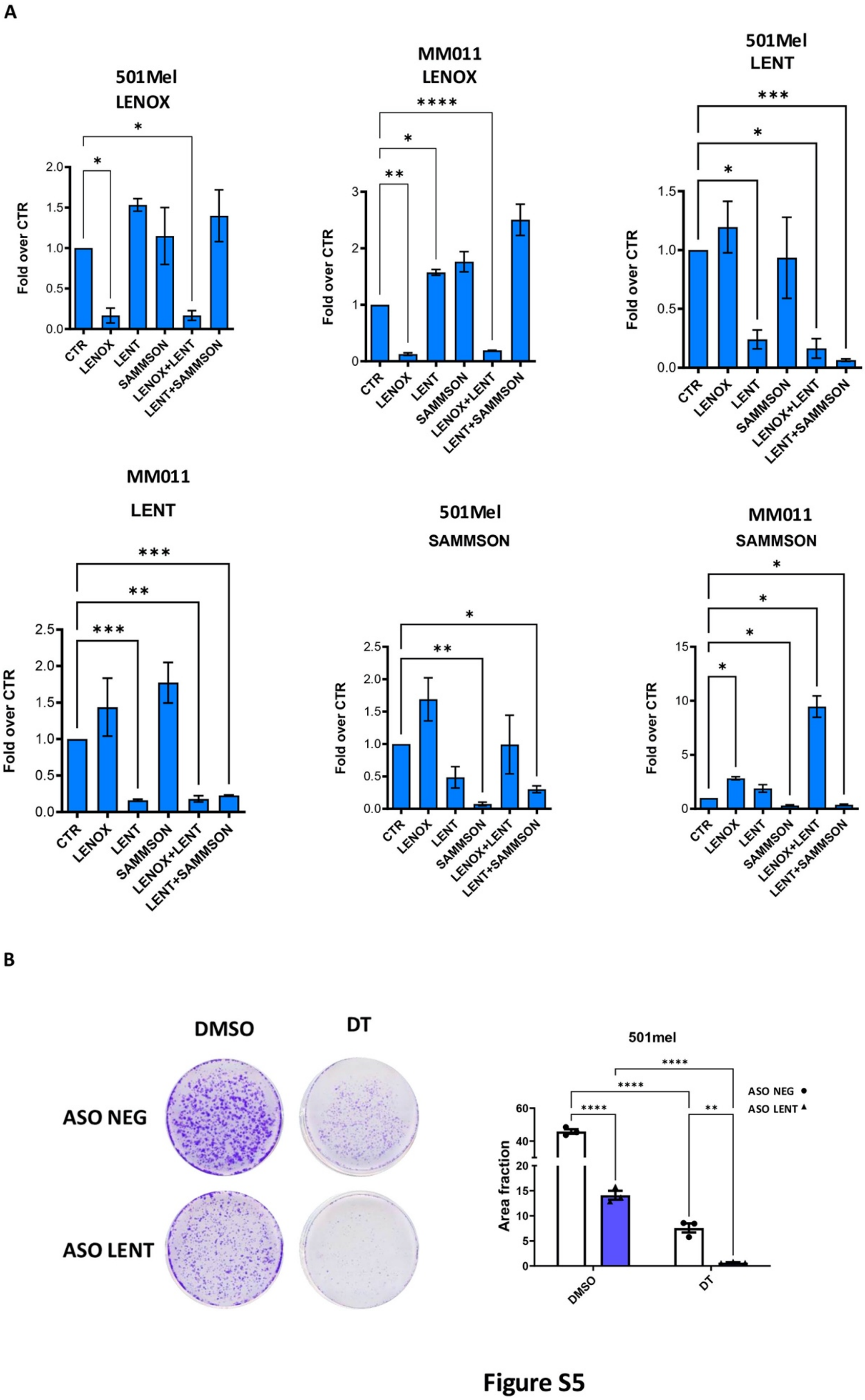
A
501Mel LENOX
MM011 LENOX
501Mel LENT
MM011 LENT
501Mel SAMMSON
MM011 SAMMSON
Fold over CTR
CTR
LENOX
LENT
SAMMSON
LENOX+LENT
LENT+SAMMSON
B
DMSO
DT
ASO NEG
ASO LENT
501mel
Area fraction
ASO NEG
ASO LENT

**Figure S5**

51

**Figure S5: LncRNA expression after combinatorial ASO knockdown. A.** Expression levels of the indicated lncRNAs measured by RT-qPCR after ASO-mediated depletion for the experiments shown in Fig 1 panels H and I. **B.** Cell proliferation was assessed after ASO-mediated LENT silencing in combination with the indicated drugs. Comparisons were done by two-way ANOVA. *, $P < 0.033$; **, $P < 0.0021$; ***, $P < 0.0002$; ****, $P < 0.0001$.

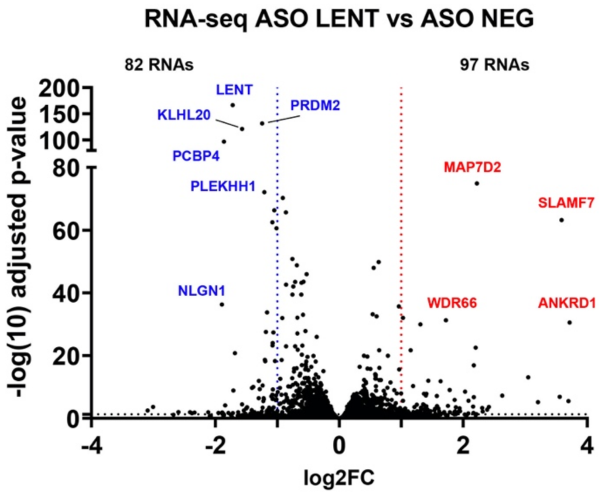


Figure S6

**Figure S6: Effect of LENT silencing on gene expression.** Volcano blot showing RNAs enriched or depleted in 501Mel cells transfected with control or LENT targeting ASO. Adjusted pvalues were derived using the Wald test.

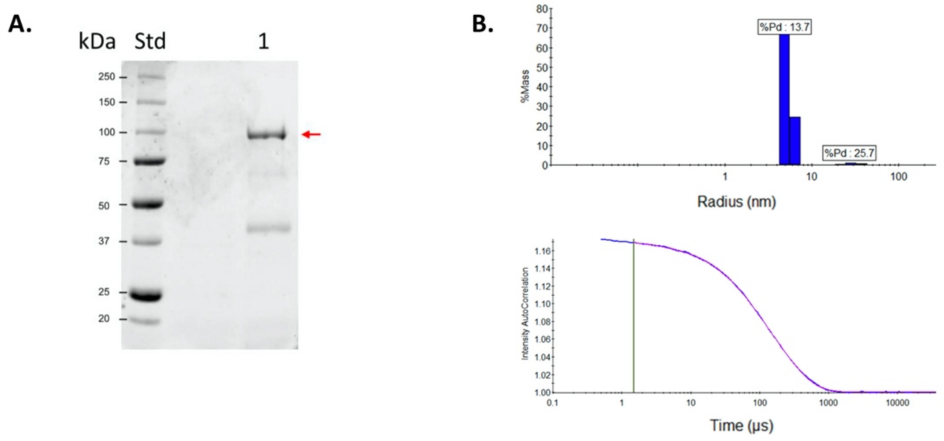


**Figure S7**

**Figure S7: DHX36 purification and LENT sequence prediction. A.** Comassie blue staining of an SDS-PAGE of purified DHX36 protein indicated by the red arrow. **B.** Upper panel, dynamic light scattering analysis of protein size distribution with Polydispersity index (%Pd) for purified DHX36; lower panel the autocorrelation function.

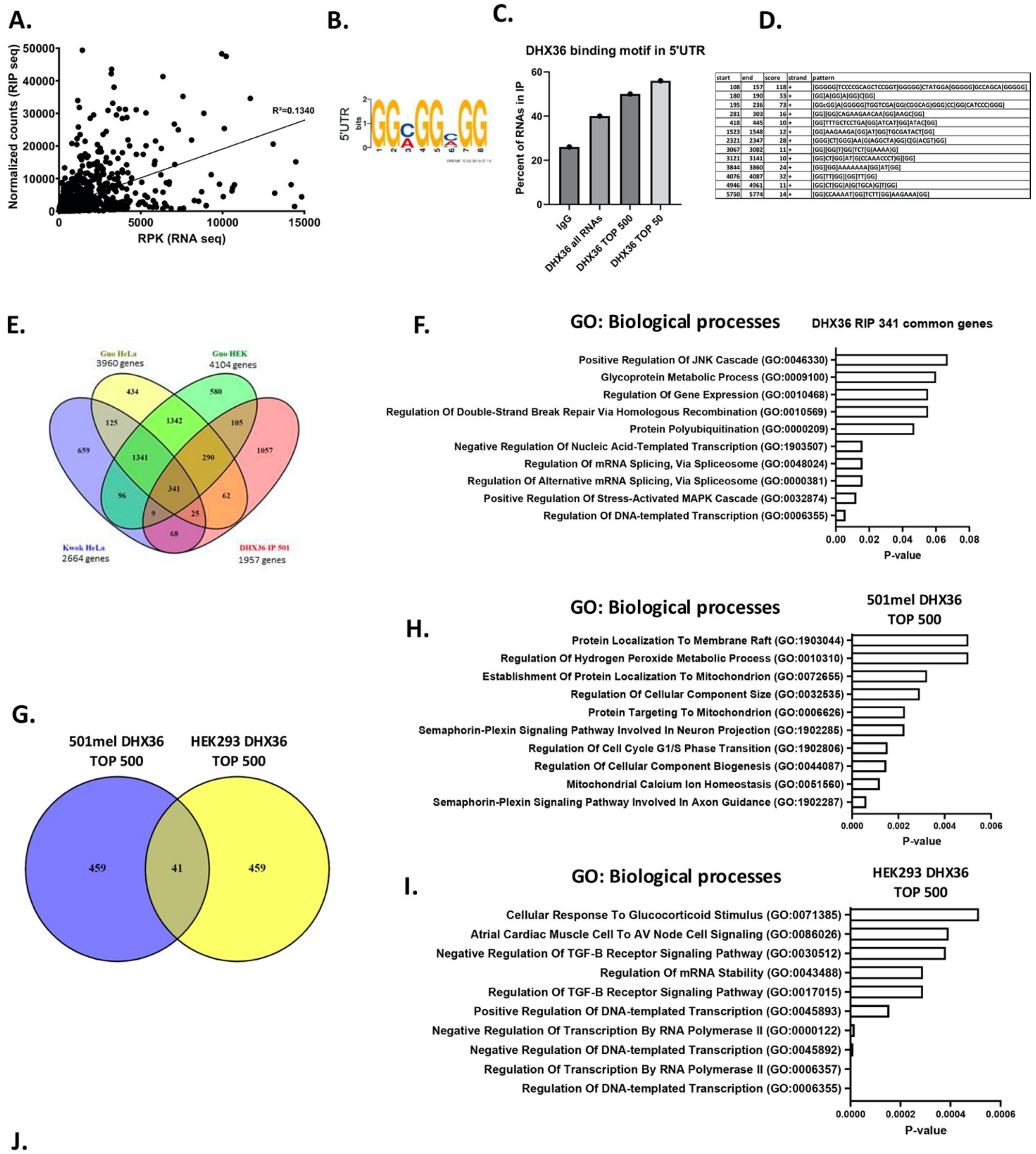


J.

| DHX36 RIP | LENT DOWN TOP 100 | LENT UP TOP 100 |
|---|---|---|
| Predicted G4 | 88/91 | 76/84 |
| Experimental G4 | 53/91 | 43/84 |
| P-value | 0 | 0 |

| DHX36 RIP | DHX36 TOP 100 | IgG TOP 100 |
|---|---|---|
| Predicted G4 | 83/91 | 6/54 |
| Experimental G4 | 71/91 | 3/54 |
| P-value | 0 | 0,2 |

**Figure S8**

66

**Figure S8. Presence of potential G4-forming structures in DHX36 associated RNAs. A.** Correlation between RNA expression level and enrichment in the DHX36 RIP calculated by linear regression. **B-C.** The % of RNAs harboring the DHX36-interaction motif shown in B in the 5'UTR of the indicated sets of RNAs was determined by the Fimo algorithm. **D.** G4 prediction by PQSfinder in the DHX36 mRNA sequence. Guanines implicated in potential G4 structures are represented in parenthesis. **E.** Venn diagram representing the overlap between RNAs enriched in the DHX36 RIP dataset reported here and the indicated data sets of potential G4-containing RNAs. 341 genes are shared between the all four datasets. **F.** Gene ontology analysis by EnrichR of the 341 common genes. **G.** Venn diagram showing the overlap between the 500 most enriched RNAs in the 501Mel DHX36 RIP seq compared to the HeLa DHX36 RIP dataset (27). **H-I.** Ontology analysis of the mRNAs specific to the 501Mel or HeLa datasets **J.** Analysis by QUADRatlas of the top 100 RNAs enriched in DHX36 RIP, control IgG RIP, or RNA whose association with DHX36 was modulated by LENT silencing. Not all genes were mapped by the algorithm explaining why the number of predicted or experimental G4s is not represented as a fraction of 100.

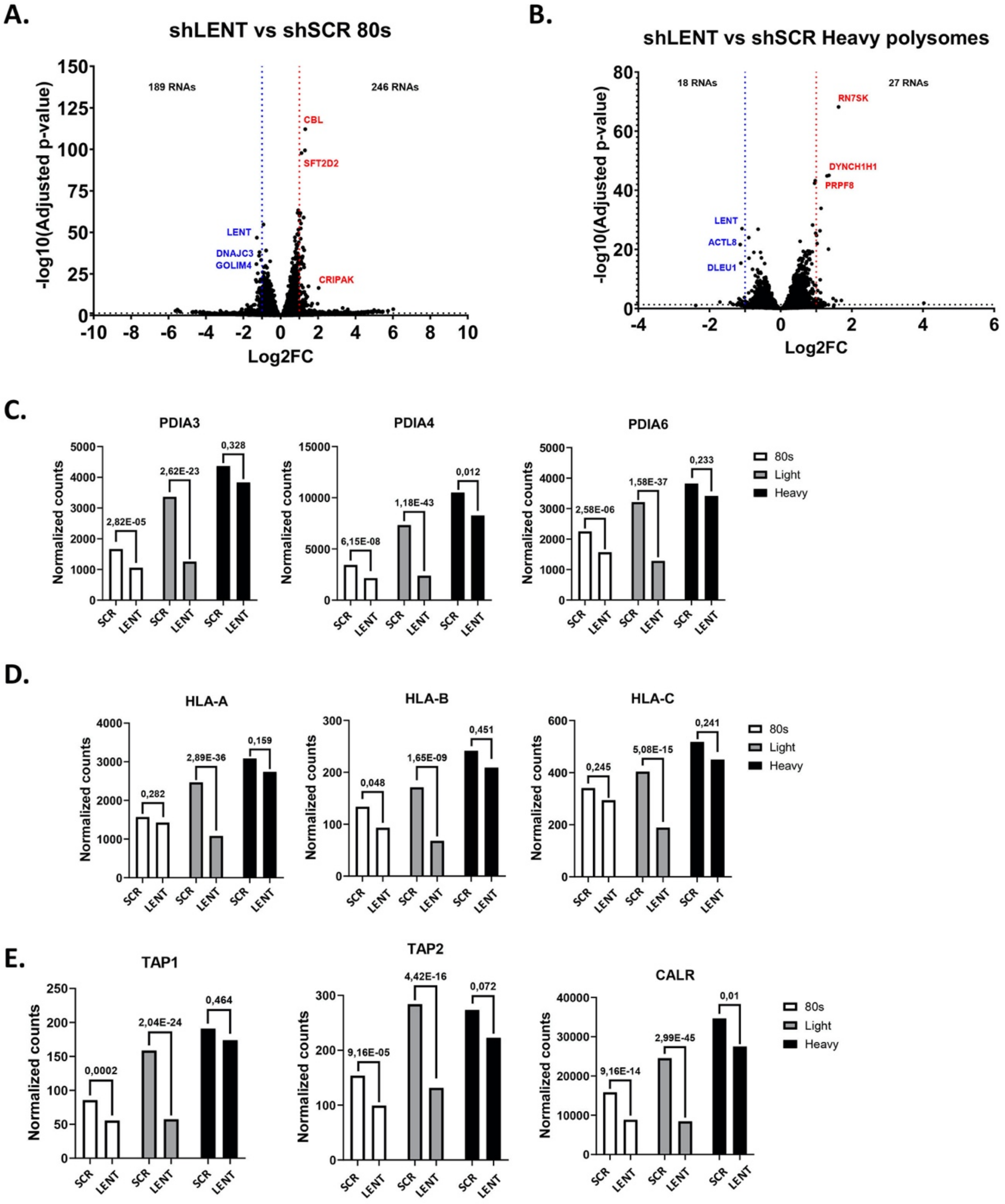


**Figure S9. Effect on LENT silencing on ribosome engagement of RNAs. A-B**. Volcano plots showing RNAs enriched or depleted in the HP and 80S fractions from the control or shLENT cells. **C-E**. RNA-seq data showing the representation of the indicated RNAs in the 80S, LP and HP fractions. The normalized number of reads are shown along with the adjusted p-value between the control and shLENT conditions.

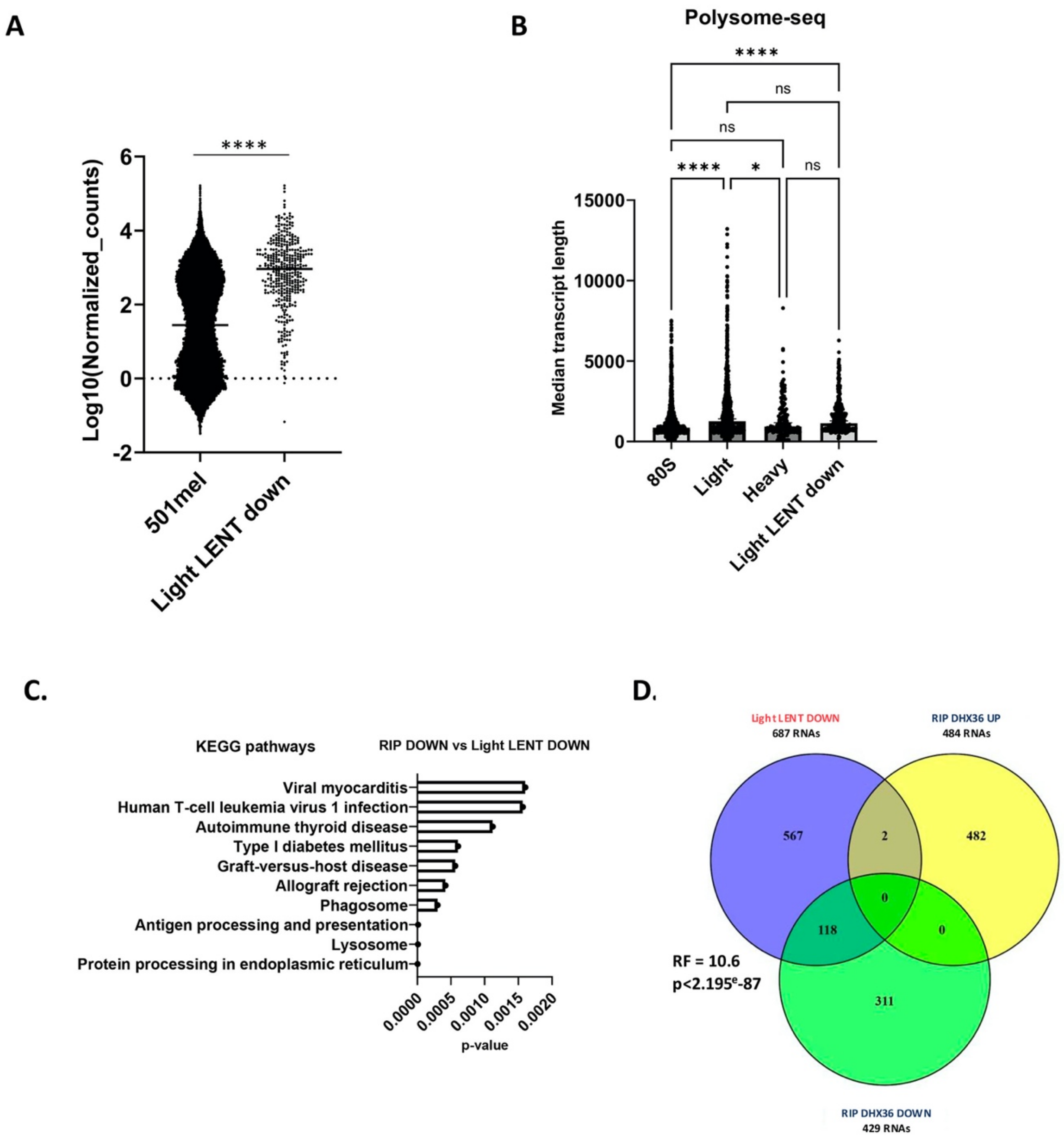


**E.**

| Light Polysomes | LENT DOWN |
|---|---|
| Predicted G4 | 342/368 |
| Experimental G4 | 217/368 |

**Figure S10.**

88

**Figure S10. Characterization of LP-depleted transcripts. A.** Expression levels of the 383 LP depleted transcripts compared to the full transcriptome. $P < 0.0001$ Mann-Whitney test. **B.** Comparison of the transcript length in the 80S, LP and HP fractions compared to the 383 LP-depleted transcripts. *, $P < 0.033$; ****, $P < 0.0001$, Kruskal-Wallis test. **C**. KEGG ontology analyses of the 74 RNAs commonly regulated by LENT in DHX36 IP and polysome profiling. **D**. Venn diagram comparing the 687 RNAs depleted in the LP fraction using the relaxed ≤ -0.8 Log2 fold change and those modulated in the DHX36 RIP in presence or absence of LENT silencing Significant Representation Factors are shown. **E.** QUADRatlas of the 383 RNAs depleted in the LP fractions.

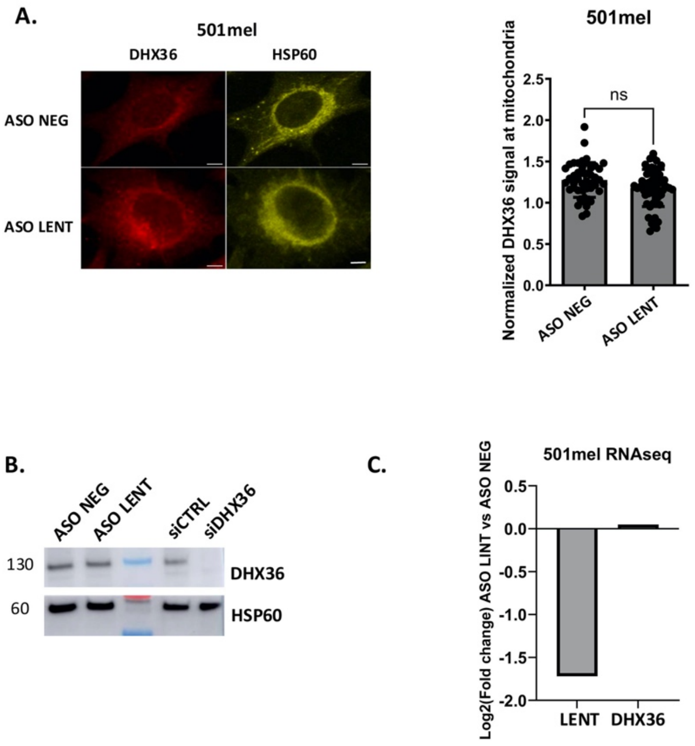


Figure S11

**Figure S11. Effects of LENT depletion on DHX36 localization or protein level. A.** Immunofluorescence by confocal microscopy showing DHX36 localization in control or LENT depleted cells. HSP60 is shown as mitochondria marker. Quantification was made by measurement of the total DHX36 signal over the DHX36 signal in mitochondria and compared to control by Mann-Whitney test. Scale bars = 2 $\mu$M. **B.** Immunoblot showing DHX36 levels upon depletion of LENT or DHX36. HSP60 used as a loading control. **C.** LENT and DHX36 RNA levels shown by RNA-seq upon LENT depletion.

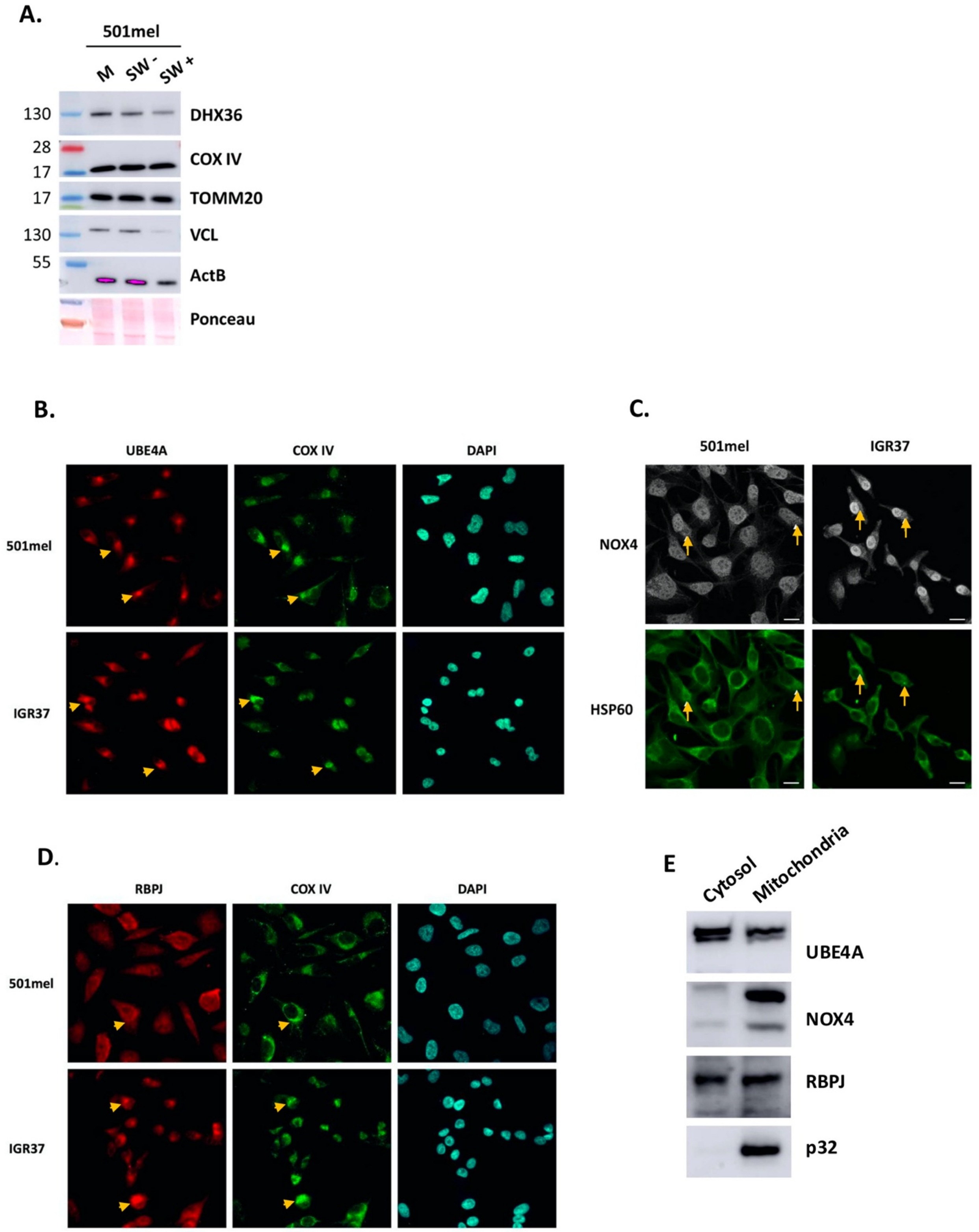


**Figure S12**

105

**Figure S12. Mitochondrial localization of proteins encoded by several mRNAs differentially bound by DHX36 upon LENT depletion. A.** Purified mitochondria were digested with trypsin with or without cell swelling buffer and remaining proteins were analyzed by western blot. **B-C.** Immunofluorescence and confocal microscopy of the indicated proteins whose mRNAs were differentially bound by DHX36 upon LENT silencing. Scale bars = 10 $\mu$M. **E.** Immunoblot after separation of mitochondria from cytosol.

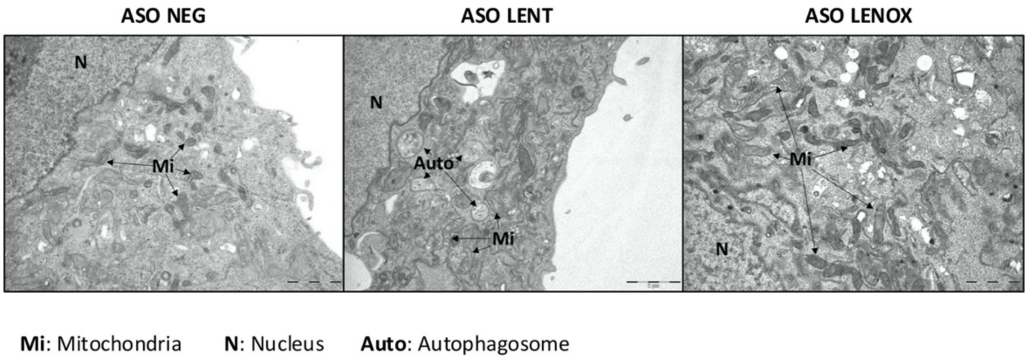


**Figure S13**

**Figure S13. Autophagy/mitophagy is specific to LENT silenced cells.** Transmission electron microscopy of 501Mel cells 48 hours following transfection of control, LENT or LENOX-targeting ASO. Scale bars are indicated on the images.

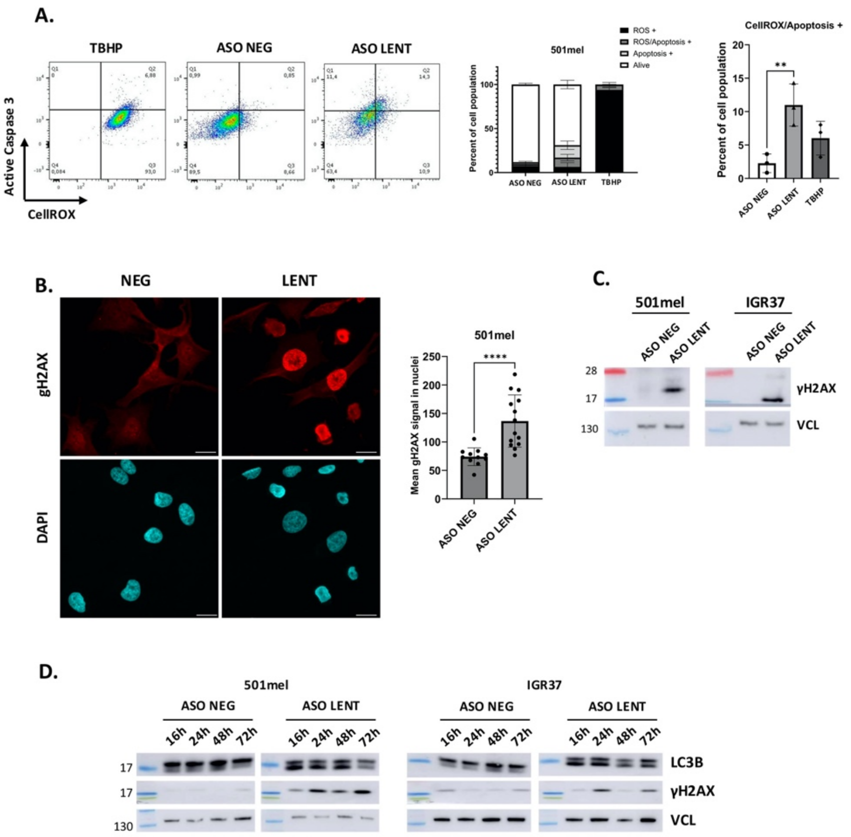


**Figure S14**

**Figure S14. ROS production and DNA damage in LENT silenced cells. A.** Flow cytometry of control or LENT-silenced 501Mel cells stained by CellROX and cleaved Caspase 3 to visualize ROS production and apoptosis. The ROS inducer TBPH was used as a positive control. Comparisons by one-way ANOVA (Dunnett test). **B.** Immunofluorescence and confocal microscopy of DNA damage marker gH2AX. Quantification of gH2AX signal in nuclei of LENT depleted cells compared to control by Mann-Whitney test. Scale bars = 10 $\mu$M. **C.** Immunblot showing gH2AX accumulation in LENT depleted cells compared to control with VCL as a loading control. **D.** Immunoblots showing accumulation of phenotypes markers at different time points upon LENT depletion. Vinculin is used as loading control.

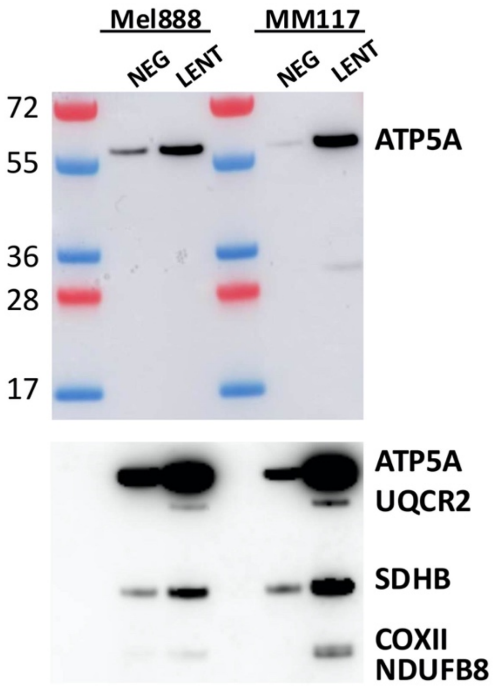


Figure S15

**Figure S15.** Immunoblots detecting mitochondrial electron transport chain proteins in extracts from Mel88 and MM117 cells. Bottom panel shows longer exposure.

**Legends to Supplemental Datasets**.

**Supplemental Dataset S1**. Excel spreadsheet showing genes up and down regulated by ASO silencing of LENT in 501Mel cells. On each page are shown gene symbol, Ensembl ID, description, gene biotype, Log2 fold-change and adjusted p-value.

**Supplemental Dataset S2**. Excel spreadsheet showing results of mass spectrometry. Shown are gene names, total number of peptides for each control (PCA3) and LENT replicate, mean values and enrichment.

**Supplemental Dataset S3**. Excel spreadsheet showing results of DHX36 immunoprecipitation. Page 1 shows RNAs enriched the DHX36 IP compared to control IgG, page 2 those depleted, page 3 RNAs that are enriched in the DHX36 IP from LENT silenced cells compared to control. Pages 4-6 show results of gene ontology of DHX36 enriched RNAs, those depleted by LENT silencing and those increased by LENT silencing corresponding to Figs. 3B and D. Pages 7-9 show the ontologies of RNAs common to the DHX36 IP and the G4 enrichment studies the 500 most enriched in the DHX36 IP from Sauer et al and the 500 most enriched in this study corresponding to Figs S6 F, H and I.

**Supplemental Dataset S4**. Excel spreadsheet showing results of RNA-seq from polysome profiling. Pages 1-6 show RNAs enriched or depleted in each fraction and page 7 the 74 RNAs common to the DHX36 IP and depleted in the LP fraction.

**Supplemental Dataset S5**. Excel spreadsheet showing on ontologies of RNAs enriched or depleted in polysome profiling. Page 1 shows biological processes of the RNAs depleted in the LP fraction, (Fig. 4G) page 2 the KEGG pathways, page 3 KEGG ontology of the common genes (Fig. S7F) and page 4 the KEGG ontology of the 80S fraction with relaxed cut off value. Page 5 shows the ontology of the transcripts with expression values >-1 and < -0.8.
